# Converging pathways found in copy number variation syndromes with high schizophrenia risk

**DOI:** 10.1101/2022.02.07.479370

**Authors:** Friederike Ehrhart, Ana Silva, Therese van Amelsvoort, Emma von Scheibler, Chris Evelo, David E.J. Linden

## Abstract

Schizophrenia genetics is complex, and the contribution of common and rare variants are not fully understood. Several specific copy number variations (CNVs) confer increased risk for schizophrenia, and the study of their effects is central to molecular models of mental illness. However, these CNVs – microdeletions or -duplications – are spread across the genome and differ in the number of genes affected and classes of coded proteins. This suggests that, in order to fully understand the contribution of these genetic variants to mental illness, we need to look beyond the deleted or duplicated genes, to their interaction partners and involved molecular pathways.

In this study, we developed machine-readable interactive pathways to enable analysis of downstream effects of genes within CNV loci and identify common pathways between CNVs with high schizophrenia risk using the WikiPathways database, and schizophrenia risk gene collections from GWAS studies and a gene-disease association database. For CNVs that are pathogenic for schizophrenia, we found overlapping pathways, including BDNF signaling, cytoskeleton, cell-cell connections, inflammation and MAPK3 signaling. Common schizophrenia risk genes identified by different studies are found in all CNV pathways but not enriched.

Our findings suggest that specific pathways – such as BDNF signaling – may be critical contributors to schizophrenia risk conferred by rare CNVs, and common risk variants may operate through distinct mechanisms. Our approach also highlights the importance of not only investigating deleted or duplicated genes within pathogenic CNV loci, but also study their direct interaction partners, which may explain pleiotropic effects of CNVs on schizophrenia risk.

## Introduction

The identification of genetic risk variants of schizophrenia has been one of the recent success stories of biological psychiatry. However, the understanding of the underlying molecular mechanisms is lagging behind. Although Genome Wide Association Studies (GWAS) identified several hundred common risk variants, each locus contributes only a small amount to the total risk (1, 2). Furthermore, many of the implicated genes were also identified as risk genes for other mental disorders with complex, multifactorial genetics and additional environmental risks factors like autism spectrum disorder, attention deficit/hyperactivity disorder, bipolar disorder and depression (1, 3). On a molecular level, the risk genes are clustered around certain biological processes and pathways that can interact with environmental events. In systems biology, a pathway is defined as a series of interactions between genes, gene products or metabolites that lead to a product or change in a cell. The Schizophrenia Working Group of the Psychiatric Genomics Consortium (PGC), identified 270 distinct loci, enriched around pathways of neuronal excitability, development and synaptic structure (4).

In addition to these associations in common variants, a number of rare copy number variants (CNVs) are recognized as highly penetrant risk factors for schizophrenia (5–7). How a CNV causes the disturbances in the metabolic and signaling network of the organism can be explained by direct gene dosage effects, unmasking of recessive variants on the unaffected chromosome (in the case of deletions) (8, 9), and downstream effects on interaction partners of the affected genes, including those caused by nuclear reorganization. Analysis of this last group of effects may give rise to the identification of affected functional pathways, some of which may overlap across CNVs.

Although these highly penetrant genetic variants can provide powerful biological models for schizophrenia and other neurodevelopmental disorders, their explanatory power and ability to yield models with high construct validity has so far been limited by a number of factors. Almost all of these CNVs include several genes, and most of these genes can be plausibly implicated in the pathophysiology of the neurodevelopmental changes (10). There is considerable locus heterogeneity - different groups of deleted or duplicated genes leading to very similar psychiatric phenotypes. The question here is what do these CNVs have in common to increase so consistently the risk of developing schizophrenia? There is also considerable phenotypic heterogeneity and variable penetrance - suggesting an important role for genetic and other modifiers (8).

Biological pathway schemes are an intuitive way to capture, understand and study such complex interactions and compare the downstream effects of different CNVs. Pathway databases like WikiPathways (11) enable the creation and modification of pathways and make them openly available for the research community. With machine-readable identifier annotations for biological entities (genes, proteins, RNA, metabolites), interactions and literature references it is possible to use these pathways for visualization and automated analysis, integration and comparison with other data resources.

In this study, we created such machine-readable pathway models that capture the genes of the schizophrenia risk CNVs and their direct interaction partners in order to find overlapping pathways or processes that might explain their increased risk and their convergence onto a unitary psychiatric phenotype. We furthermore investigated if these pathways host more known schizophrenia risk genes than expected by average distribution across the genome.

## Results

### 1. Pathway creation

We created pathway models for the defined CNV syndromes in order to provide the available interaction knowledge of the affected genes in a machine-readable way that enables further analysis. These pathways are available through the rare disease portal of WikiPathways http://wikipathways.org. Table 1 summarizes the pathways and their main characteristics. Figure 1 shows an example of one of these pathways. For about 10% of the protein coding genes in these affected regions, there was no information on their function or interactions available, yet.

**Table 1:**
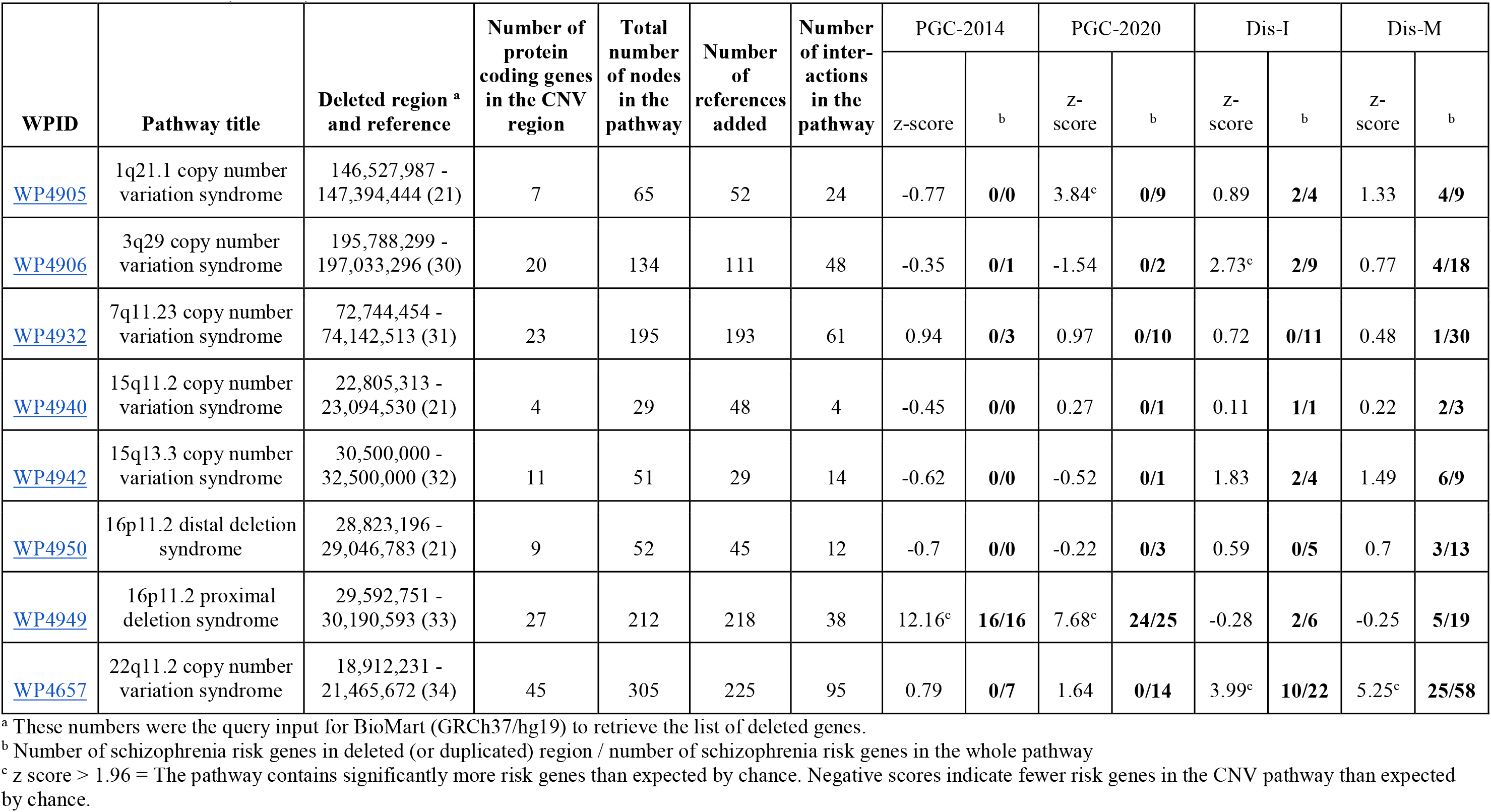
CNV pathways created in this study and presentation of schizophrenia risk genes from different sources in those pathways by number and enrichment score (z-score).

**Figure 1:**
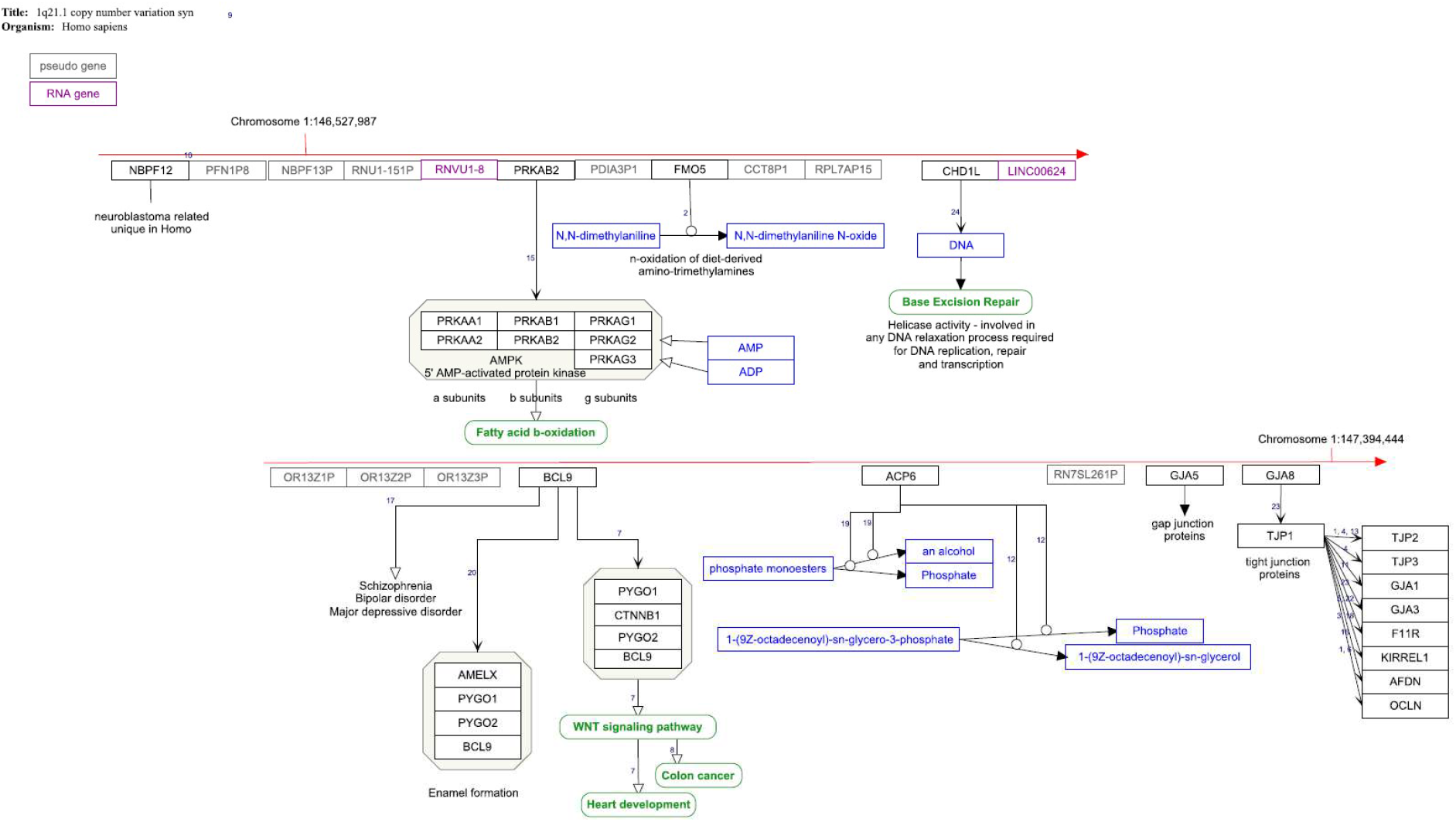
1q21.1. Copy number variation pathway WP4905. The red arrow indicates the position of the deleted genes on the chromosome. Protein coding gene are black boxes, pseudogenes grey, RNA genes purple, metabolites blue and pathways green. The interactions are annotated interactions carrying the information about the nature of interaction for automated analysis. The small numbers indicate the literature references http://www.wikipathways.org/instance/WP4905.

### 2. CNV pathway overlap analysis

To identify potential overlap of the different CNV pathway models that might explain common observed symptoms, the pathways were imported to Cytoscape, extended with gene-pathway information from WikiPathways database adding known pathways to a gene, forming a gene-pathway network. Pathways that shared genes with more than one CNV were extracted. All CNVs shared genes from the “Brain-Derived Neurotrophic Factor (BDNF) signaling pathway”. If the smallest pathway (15q11.2, four protein coding genes) was removed, there were further seven pathways (out of a total of 1453 pathways approved for data analysis in the database) which shared genes with the remaining CNV pathways: “Ciliary landscape”, “Leptin signalling pathway”, “VEGFA-VEGFR2 Signalling Pathway”, “TNF related weak inducer of apoptosis (TWEAK) Signalling Pathway”, “IL-18 signalling pathway”, “Focal Adhesion”, and “Gastrin Signalling Pathway” (Table 2). Five of eight CNVs (not 15q11.2, 16p11.2 distal, and 22q11.2) shared genes with at least one, but usually several, WNT signaling pathways.

**Table 2:** Shared pathways between the different CNVs and their z-scores from schizophrenia risk gene overrepresentation analysis.

| Pathway | ID | PGC-2014 | PCG-2020 | Dis-I | Dis-M |
| --- | --- | --- | --- | --- | --- |
| Brain-Derived Neurotrophic Factor (BDNF) signaling pathway | WP2380 | 1.89 | 1.39 | 6.65<sup>a</sup> | 7.31<sup>a</sup> |
| Ciliary landscape | WP4352 | -0.62<sup>b</sup> | -1.32<sup>b</sup> | -3.84<sup>b</sup> | -2.01<sup>b</sup> |
| Leptin signaling pathway | WP2034 | -0.41 | 0.19 | 3.34<sup>a</sup> | 2.51<sup>a</sup> |
| VEGFA-VEGFR2 Signaling Pathway | WP3888 | 0.89 | 1.24 | 2.84<sup>a</sup> | 1.77 |
| TNF related weak inducer of apoptosis (TWEAK) Signaling Pathway | WP2036 | 0.19 | 0.22 | 2.32<sup>a</sup> | 2.27<sup>a</sup> |
| IL-18 signaling pathway | WP4754 | -1.04<sup>b</sup> | -1.36<sup>b</sup> | 2.53<sup>a</sup> | 1.2 |
| Focal Adhesion | WP306 | -0.98<sup>b</sup> | -0.23<sup>b</sup> | 1.81 | 0.82 |
| Gastrin Signaling Pathway | WP4659 | 1.85 | 2.09<sup>a</sup> | 4.51<sup>a</sup> | 3.84<sup>a</sup> |

<sup>a</sup> z-score > 1.96 indicates a significantly overrepresented pathway, indicating more schizophrenia risk genes than expected by chance.
<sup>b</sup> z-score < 0 indicates a pathway with fewer schizophrenia risk genes than expected by chance. < -1.96 indicates significantly fewer.

In a second step, the extended CNV pathways were merged, and network analysis revealed the most densely connected nodes within the network indicating hubs of importance (Supplementary Table 2). The larger CNV pathways (without 15q11.2 and 15q13.3) were among the original top ten but were removed from the list in order to avoid redundancy. The full network is available on Ndex database [https://doi.org/10.18119/N96W3R].

The links between pathways formed by shared genes are defined by prior knowledge from experimental evidence as stated in the literature references of the individual pathways. However, to investigate how likely it is that a random gene set shares genes with the pathways listed in Table 2, permutation statistics were done (Supplementary Information 2). In short, it is highly unlikely that the connection patterns we found could have occurred by chance. The number of randomly overlapping genes is, as expected, strongly dependent on the size of the pathways. For example, an average of 3.4 ± 1.2 of eight pathways share a random gene with the BDNF pathway (which has 144 genes) by chance. The actual overlap was much higher also for the other pathways investigated. The VEGFA-VEGFR2 signaling pathway, with 431 genes the largest pathway in this study, was an exception. It showed a high random overlap with the CNV pathways due to its size.

To investigate the benefit of downstream interaction partners to the genes affected by CNVs another network was created using only the genes deleted/duplicated in these CNVs. In this approach, no overlapping pathways between the different CNVs were found (data not shown).

The genes, which form the connections between the CNV pathways and the eight pathways (Table 2) were isolated and visualized in Figure 2a. As the BDNF pathway is of special interest in brain function and development, the network connecting it with the CNV pathways is shown in detail in Figure 2b. In the BDNF signaling pathway network (Figure 2b), most genes connect to one of the CNV pathways. There are three genes, which are present in more than one CNV pathway: CTNNB1, GRB2, and FYN. CTNNB1 is also one of the most connected hub genes in the network.

**Figure 2:**
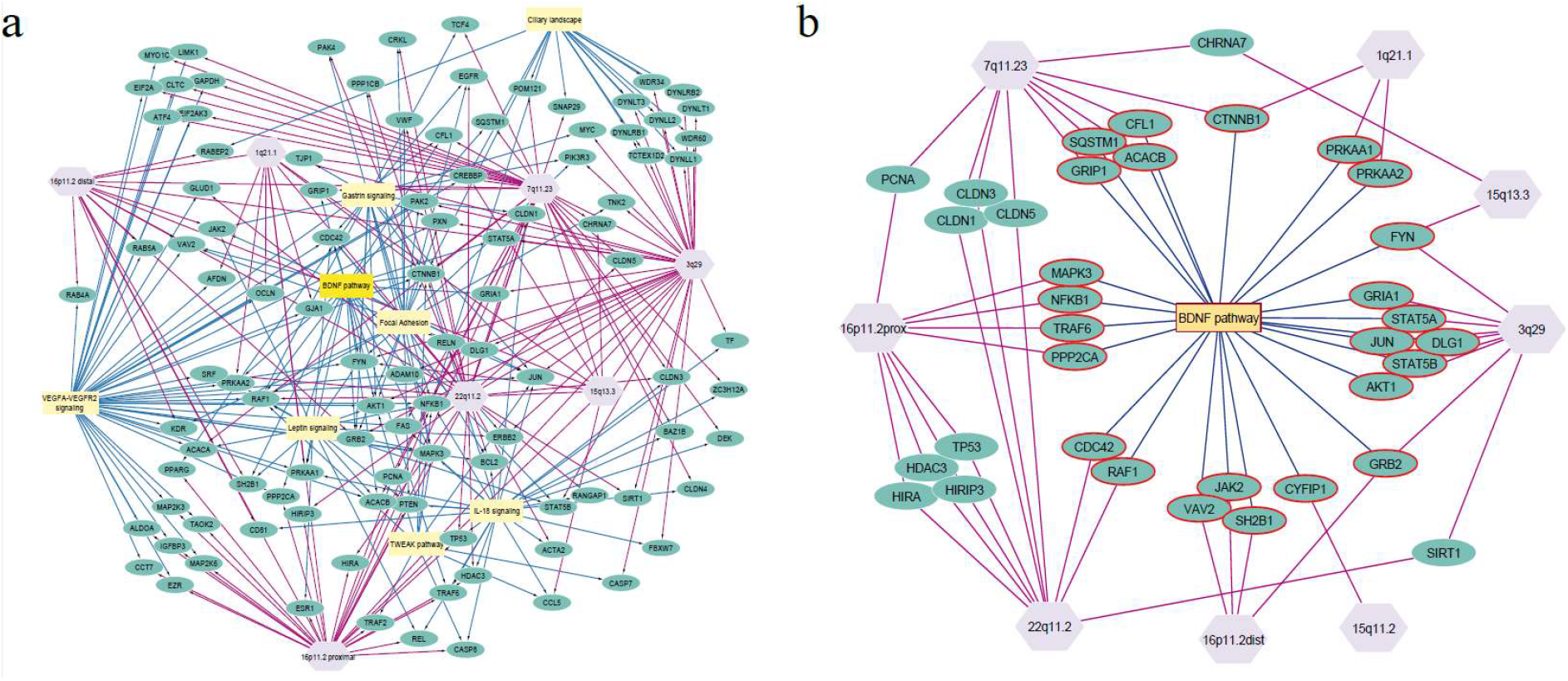
Pathway overlap between the different CNVs. a) The seven CNV pathways (violet octagons) (without 15q11.2) share genes (green spheres) with a group of pathways (yellow rectangles, list in Table 2) that are known to play a role in schizophrenia development. Edges from the CNV pathways are marked in red, edges from connected pathways in blue. b) BDNF pathway shares between one to six genes (red) with each of the investigated CNV pathways.

### 3. Comparison with known schizophrenia risk genes

There were 331 (PGC-2014) schizophrenia risk genes from the earlier and 3 627 (PGC-2020) from the most current publication of the PGCs GWAS extracted and mapped to Ensembl identifiers. The DisGeNET variant-disease association list for schizophrenia contained 2 897 unique variants. 2 330 variants were mapped to 1 392 different protein-coding genes. The gene-disease-association list contained 2872 genes. These two DisGeNET lists share 949 genes (intersection, Dis-I). The merged list of both, gene and variant-disease associations, contains 2701 genes (merged, Dis-M).

There is some overlap between the different datasets (Figure 3). DisGeNET includes gene disease and gene variant information *i.a*. from the NHGRI-EBI GWAS Catalog where the PGC-2014 study is listed. The number of genes listed there is lower than given by the original authors because DisGeNET itself filters out variants with p>1.0×10^-6^ and a predefined group of variant consequences. The most recent PGC study is not listed in the catalog, yet.

**Figure 3:**
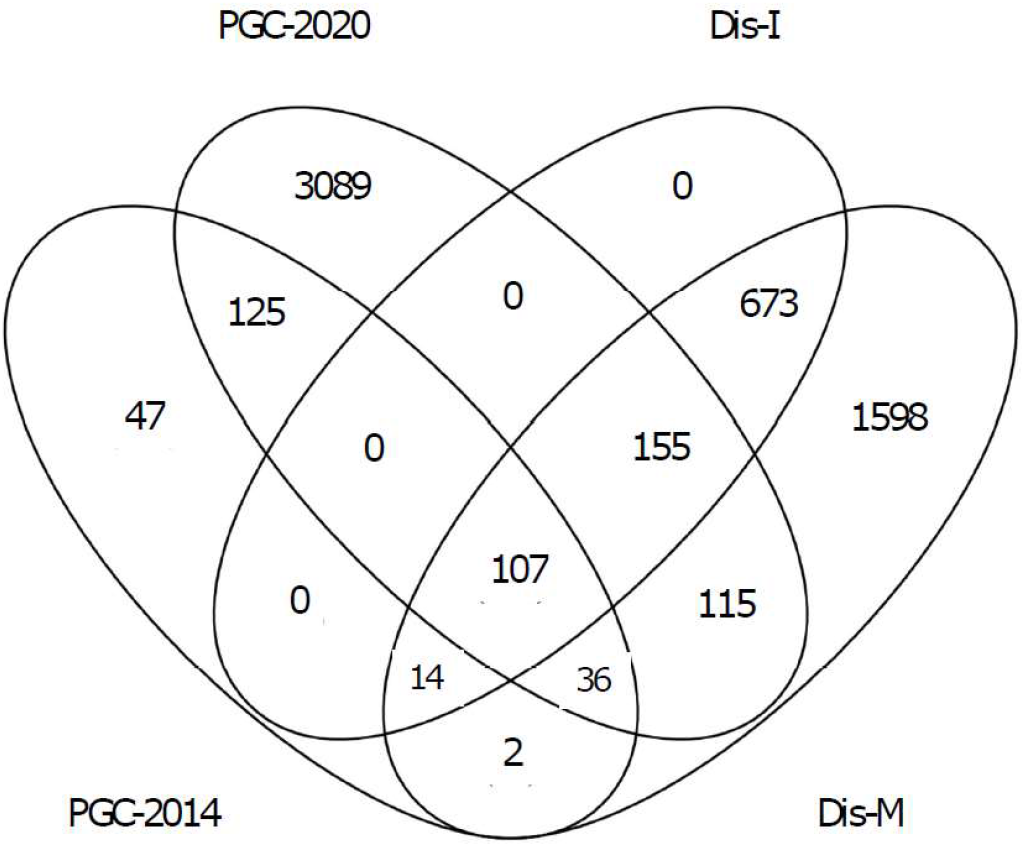
Overlap in schizophrenia risk genes extracted from GWAS studies and DisGeNET database. PGC-2014 (2) and PGC-2020 (4) from their respective publications, Dis-I = intersection/overlap between DisGeNET’s gene-disease and variant-disease associated genes for schizophrenia, Dis-M = merge of DisGeNET’s gene-disease and variant-disease associated genes for schizophrenia.

#### Visualization and overrepresentation of schizophrenia risk genes in CNV pathways

Table 1 also shows the number of schizophrenia risk genes in the different CNV pathways. We indicated how many risk genes are located specifically in the affected CNV locus and how many are found in the whole pathway (Supplementary pathway figures). Enrichment analysis z-scores above 1.96 show that these pathways host more than the average number of schizophrenia risk genes expected by chance for this dataset.

The PGC datasets identified a part of the 16p11.2 proximal locus as a risk locus. PGC-2014 identified 16 of the 27 genes on this locus, and PGC-2020 identified 23. Consequently, the whole pathway got a high enrichment score. Similarly, for 1q21.1, there were nine consecutive risk genes found. For this, the whole pathways got a high enrichment score. Notably, apart from the 1q21.1 and the 16p11.2 proximal locus, there are hardly any risk genes identified by the PGC datasets that are in the regions of the other CNV pathways, although there was enrichment for several downstream genes, e.g. 15 genes of the 22q11.2 and 10 genes in the 7q11.23 pathway (Table 1).

Schizophrenia-associated genes from the DisGeNET dataset are generally found more often in the CNV pathways. As the PGC-2020 (3 627) and the Dis-M (2 701) dataset are of comparable size we presume that this is not primarily an effect of dataset size.

#### Pathway enrichment results

To investigate which other than CNV pathways contained an increased number of schizophrenia risk genes a pathway overrepresentation analysis was done using the different risk gene lists and the WikiPathways database (full results in Supplementary Table 3). For all lists, the “Alzheimer’s disease”, “Prion disease pathway”, “Rett syndrome causing genes” and “Serotonin receptor 4/6/7 and NR3C signaling pathway” were enriched. Common in both PGC and DisGeNET datasets were further: pathways around neurological function (e.g. “Nicotine Activity on Dopaminergic Neurons”, “PKC-gamma calcium signalling pathway in ataxia”, “Disruption of postsynaptic signalling by CNV”), lipid metabolism (“Lipid homeostasis”, “Arachidonate Epoxygenase/Epoxide Hydrolase”), immune system (“Complement and Coagulation Cascades”), WNT signaling (“Regulation of Wnt/B-catenin Signalling by Small Molecule Compounds”) and “Genes involved in male infertility”. Notably, most of the overlapping pathways between the different CNVs are enriched with at least one schizophrenia risk gene list (Table 2). The pathways associated with cytoskeleton and cilia function have a negative z-score, indicating a smaller number of risk genes than expected by chance.

## Discussion

### 1. Resource creation

In this study, we created pathway models for several CNVs with a high penetrance for schizophrenia, which serve as a bioinformatics and data analysis resource to enable pathway and network modelling and analysis of any kind of omics data. The pathways are fully machine-readable, annotated with unique, persistent identifiers from resources like Ensembl, UniProt or ChEBI, and provide the provenance for every single statement in form of literature references. The pathways can be found in the WikiPathways database and will undergo regular curation and updates (11).

### 2. Overlapping pathways analysis

Using the information about downstream interaction partners of the deleted/duplicated genes in the different CNVs it was possible to identify pathways on which these CNVs converge (Table 2). These shared pathways align with previously known pathways and biological processes affected in schizophrenia:

1. Neuronal development and function, as seen in the BDNF pathway. All investigated CNV pathways have direct links to BDNF-triggered downstream functions (Figure 2b) (12) but also to other processes that are central neuronal development. For example, CYFIP1 (15q11.2 locus) is found in the Fragile X syndrome pathway, interacts in the wave regulatory complex to regulate cytoskeleton remodeling by regulating the production of f-actin, and is generally important for brain development (13, 14).
2. Cytoskeleton and cell-cell connections, including ciliary landscape, gap junction (note that two gap junction proteins (GJA5, GJA8) are located in the 1q21 region and associated with cardiac and ocular phenotypes (15)) and focal adhesion pathways. These contain proteins that are also involved in synaptic structure, e.g. KCTD13 (16p11.2 prox locus) (16). In addition to its role in axonal growth, CTNNB1, which is part of the 1q21.1 and 7q11.23 pathways, can also be found in the Ciliary landscape pathway (17).
3. Immune system pathways are also highly affected and connected within all CNV pathways, specifically, the TNF related weak inducer of apoptosis (TWEAK) and the IL-18 signaling pathway. The TWEAK pathway has recently been identified a core pathway in depression (18).

Common signaling hubs, like the PI3K-Akt Signaling Pathway and the MAPK3 pathway, were identified both as a direct overlapping pathway between all CNVs. *MAPK3* is located at 16p11.2 and therefore directly affected by deletion/duplication but due to its broad action range it also affects almost all other CNV pathways (19). Generally, MAPK3 and several other kinases and transcription factors such as AKT1, NKFB1, JUN and TP53 tend to show up as network hubs because they are so frequently studied and involved in so many different processes. Thus, their involvement in the CNV pathways may not be particularly specific.

*CTNNB1* is a highly connected hub gene in the network and it is one of the genes connecting two CNV pathways (1q21.1 and 7q11.23) and the BDNF signaling pathway (Supplementary Table 3, Figure 2b). In the 1q21.1 pathway it is a binding partner in a complex with BCL9 (whose gene is located in 1q21.1), PYGO1 and PYGO2, contributing to the WNT signaling pathway. In the 7q11.23 CNV pathway it is inhibited by BCL7B (whose gene is located in the 7q11.23 region), which is therefore also an inhibitor of the WNT signaling pathway. WikiPathways hosts several individual WNT signaling pathways that cover different aspects of up- or downstream effects. Most CNVs share a link to at least one of those WNT signaling pathways, which makes it an interesting pathway to study for these disorders.

To conclude, using the CNV pathways, it was possible to identify common processes shared by all these CNVs with high risk of developing schizophrenia share, which may lead to new methods for targeting these pathways for early diagnosis or treatment.

### 3. Investigation of schizophrenia risk gene distribution and identification in the CNV pathways

The identification of these CNVs and their shared pathways invites the question of genetic links between these (relatively) rare forms of schizophrenia and the large number of common variants identified by GWAS. One could hypothesize that these pathways host a number of schizophrenia risk genes, higher than expected by average. To answer this question we conducted overrepresentation analysis and chose two different collections of schizophrenia risk genes: The first datasets come from GWAS studies PGC-2014 (2) and PGC-2020 (4) and another from a gene-disease association database DisGeNET (20).

Although using different methods for functional enrichment analysis, the result of the pathway analysis of the GWAS study (2), “neuronal excitability, development, and structure, with prominent enrichment at the synapse”, matches with our results for the CNV pathways: post-synapse location, neuron projection, and synaptic signaling.

The DisGeNET database collects a broader spectrum of gene-disease and variant-disease associations by integrating curated databases, animal model data, inferred data and literature (20). In contrast to GWAS data, DisGeNET includes both common and rare variants in the gene-disease associations.

Visualizing the schizophrenia risk genes in the CNV pathways showed clearly that all pathways host schizophrenia risk genes (except the smallest risk gene list of PGC-2014). These risk genes are not always located in the deleted region but there are always some direct interaction partners of the deleted genes, which are known to be a risk gene. Given the about 3 000 potential schizophrenia risk genes (more than 10% of the genome!) this is not surprising.

The question is, whether there is an unusual accumulation of risk genes in these pathways. This was answered in this study using overrepresentation analysis. To summarize, this analysis showed that except for 3q29 and 22q11.2 (rare and common variants) and 1q21.1 and 16p11.2 prox (common variants), the CNV pathways are **not** hosting more than the expected average number of schizophrenia risk genes – whereas the majority of the overlapping pathways between the CNVs does show such enrichment for schizophrenia risk genes. 3q29 and 22q11.2 CNV pathways are highly enriched for the DisGeNET datasets and are also the CNVs with the highest known schizophrenia risk (5, 21). 1q21.1 and 16p11.2 was itself identified in the PGC GWAS studies as a high-risk locus and is therefore, not surprisingly enriched in this analysis.

The conclusion from this study emphasizes that the increased schizophrenia risk in certain CNVs originates from both the quantity of affected risk genes in the CNV region itself (as shown for 16p11.2, 22q11.2 and 3.29) and the connection to certain biological pathway processes. The effects of CNVs on pathways related to BDNF signaling, cytoskeleton, and immune system, and also lipid metabolism (22) and WNT signaling, could constitute relevant pathophysiological mechanisms.

### 4. Strengths and Limitations

The focus of this study is not on single genes but on whole pathways, supporting the emerging pathway/network based medicine and creating a systems biology resource for future investigations of the genetic mechanisms of schizophrenia and other neurodevelopmental disorders.

Pathway creation is dependent on literature input (with implicit bias), requires time and experience to map biomedical knowledge properly to machine-readable identifiers and interactions. We cannot claim completeness of the pathway in reference to the known interaction partners of the genes in the CNV loci investigated. However, WikiPathways is a community-created, expert-curated pathway database, and thus, future improvements and updates can easily be incorporated. A limitation of our overlap analysis with DisGeNET is that this database incorporates information from CNV studies, but with the low number of genes in the CNV loci compared to the overall number of schizophrenia risk genes listed in DisGeNET the influence of such circularity on our results would have been minimal.

### 5. Conclusion and Implications

We identified several potential underlying pathways, most importantly BDNF signaling, that connect pathogenic CNVs on a molecular level and might explain their shared high penetrance for schizophrenia. A key implication of this work relates to the improvement of our understanding of convergent mechanisms of genetic risk for schizophrenia, and further work should determine whether similar mechanisms could also be identified in schizophrenia patients without high-penetrance variants. Another line of impact is the contribution to drug discovery, based on the identification of drug targets in the nodes of the identified pathways. Further refinement of this work might concern the separation of developmental stages in the pathways, which may enable a focus on the early and adolescent brain development that is so relevant for the genesis of schizophrenia.

## Methods

### CNV selection

The CNV loci with an increased risk of psychiatric disorders (through deletion, duplication or both) were selected as proposed by Kirov et al. (5) and include 1q21.1, 3q29, 7q11.23, 15q11.2, 15q13.3, 16p11.2, and 22q11.2.

### Pathway construction

To construct the pathways, PathVisio pathway drawing software was used (23) including gene product and metabolite mapping databases from BridgeDb (24). First, for each CNV, a list of genes located in the typically deleted or duplicated region was queried from Ensembl genome browser, Human Genome Build GRCh37/hg19 using BioMart (25). The exact genome positions are summarized with the respective references in Table 1. The genes were imported to PathVisio and different sources, such as scientific literature and databases, were investigated to find information, reactions, interaction partners and downstream pathways for the protein coding genes. A detailed description of the search strategy is given in Supplementary Information 1.

The nodes in the pathway, such as genes, proteins, RNA and metabolites, were annotated with Ensembl (genes and gene products) or ChEBI (metabolites) identifiers. Interactions were annotated with MIM identifiers, which indicate the nature of interaction and, if available, with their respective RHEA identifiers [www.rhea-db.org]. The interactions also bear the identifier of the source, usually a PubMed identifier of the study that describes the interaction.

### Network construction

Cytoscape (26) and associated packages (WikiPathways (27), RCy3 (28)) were used to import the new CNV pathways to Cytoscape, extend them with pathway information (CyTargetLinker (29)), merge, and analyse them. The WikiPathways linksets wikipathways-hsa-curated-20210110 and wikipathways-hsa-Reactome-20210110 were used.

### Schizophrenia risk genes

Lists of known schizophrenia risk genes were acquired from two different resources. First, we identified publications with lists of common schizophrenia-associated loci, identified by GWAS. A list of 349 associated genes (of which 331 could be mapped to a gene identifier) was extracted from the table of 108 risk loci published by the Schizophrenia working group of the PGC (2), extracted from their Supplementary Table 3 (PGC-2014). A second list was extracted from this consortiums’ most recent publication (4), also their Supplementary Table 3, resulting in 3 627 genes (PGC-2020).

Second, we consulted the DisGeNET database (20) [Dec 1st, 2020] for gene-disease and variant-disease associations from multiple curated resources. We used the DisGeNET search function to extract genes and genetic variants associated with schizophrenia [CUI: C0036341]. The variants were mapped to genes before further use.

All gene lists are available in Supplementary Table 1.

### Overrepresentation analysis

Pathway overrepresentation analysis was done using PathVisio statistics function described in (23). This is based on a commonly used overrepresentation statistics, calculating the z-score, which gives an indication if there are more genes from a chosen dataset – in this case, the risk gene lists - found in a specific pathway, compared to the number of genes expected by chance. The background list was a standard list of protein coding genes from Ensembl containing about 20 000 human genes. The pathway collection was downloaded from WikiPathways database (version 20201110).

## Supporting information

Supplementary CNV pathway figures

Supplementary overlapping pathways figures

## Acknowledgements and funding sources

The authors would like to thank Prof. Dr. Han Brunner for helpful discussions around genetics of neurodevelopmental disorders, and Dr. Martina Kutmon and Dr. Lars Eijssen for helpful discussions around permutation testing statistics. FE and CE are funded by the European Union’s Horizon 2020 research and innovation programme under the EJP RD COFUND-EJP N° 825575. TvA is funded by NIH_5U01 MH119740. The authors declare no conflict of interest.

