## Supplementary CNV pathway figures for "Converging pathways found in copy number variation syndromes with high schizophrenia risk"

**Title:** 1q21.1 copy number variation syn

**Organism:** Homo sapiens

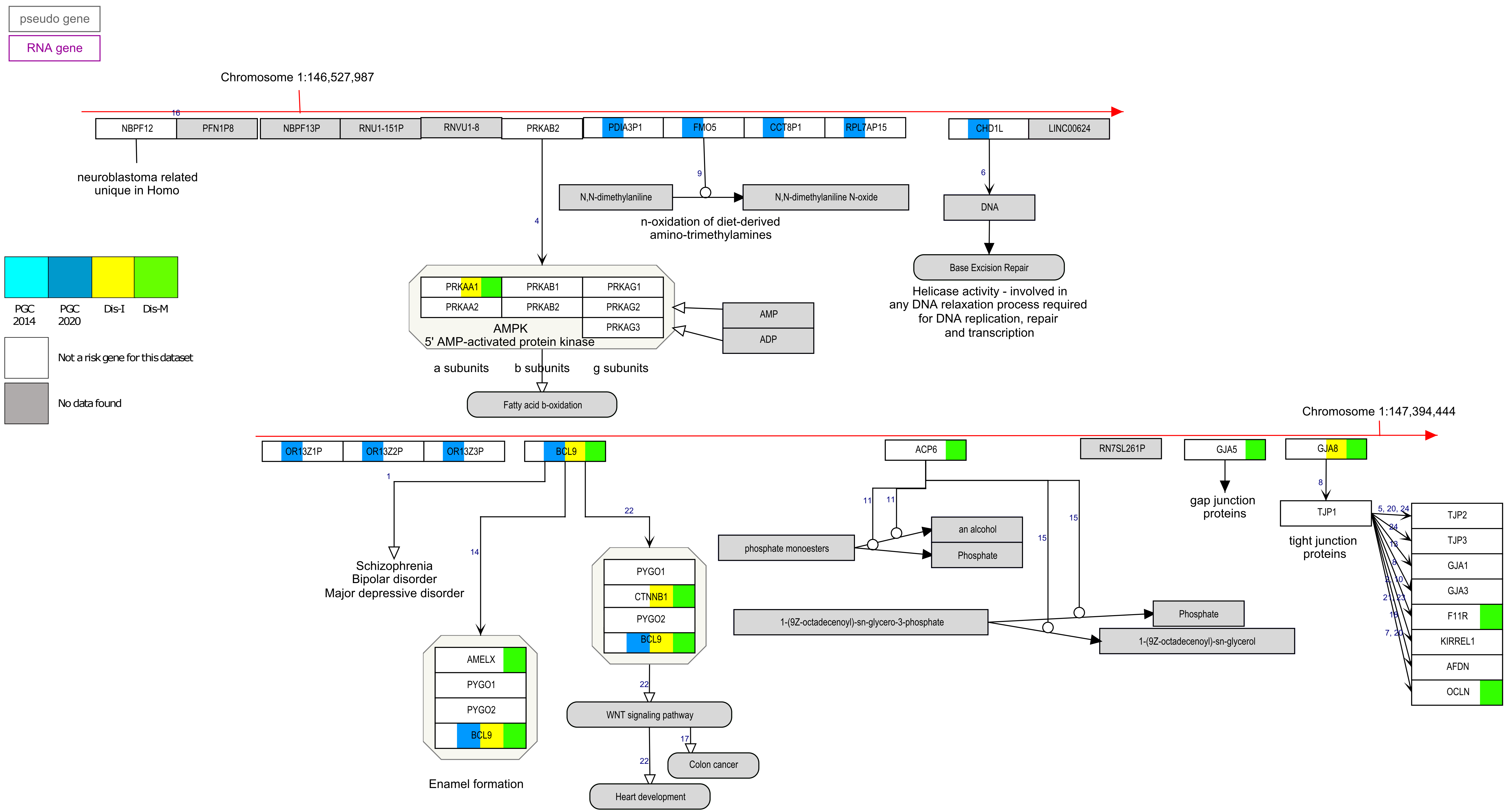

**Title:** 3q29 copy number variation syndr
**Organism:** Homo sapiens

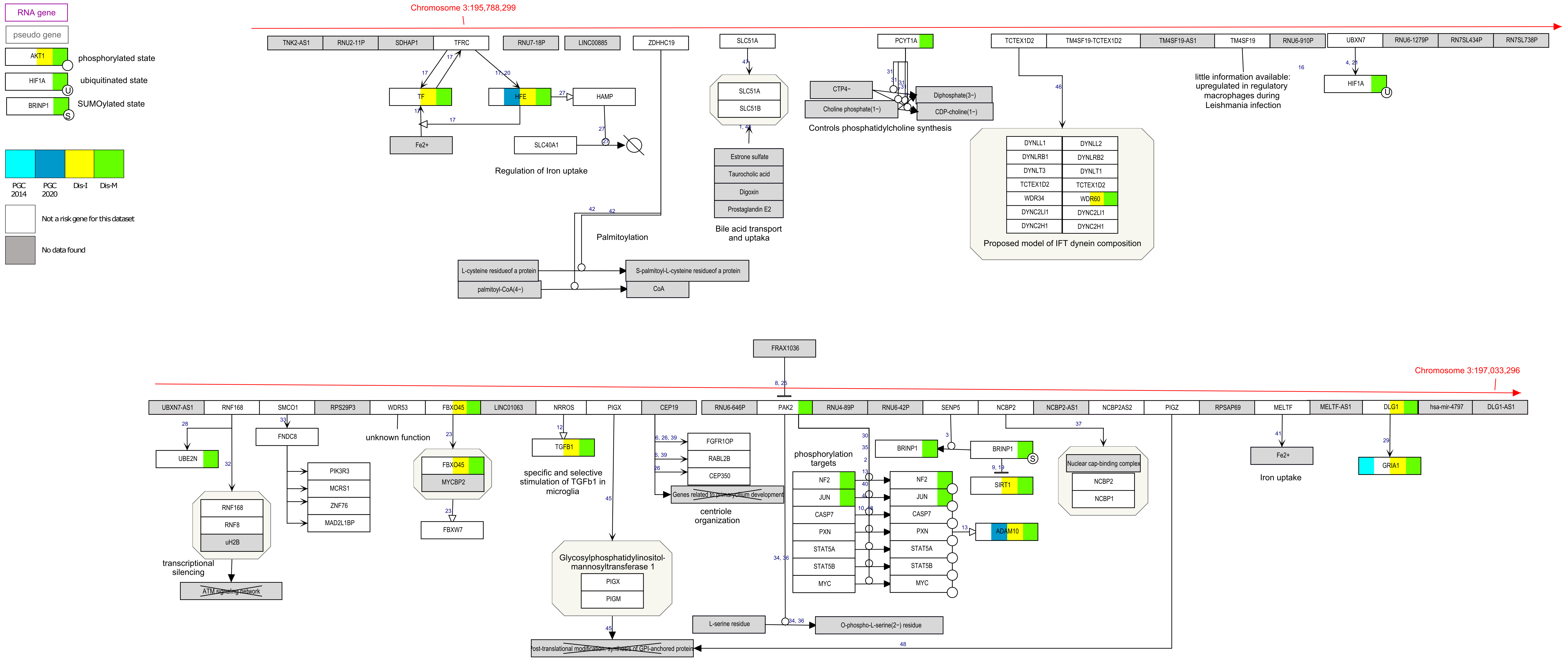

**Title:** 7q11.23 copy number variation sy
**Organism:** Homo sapiens

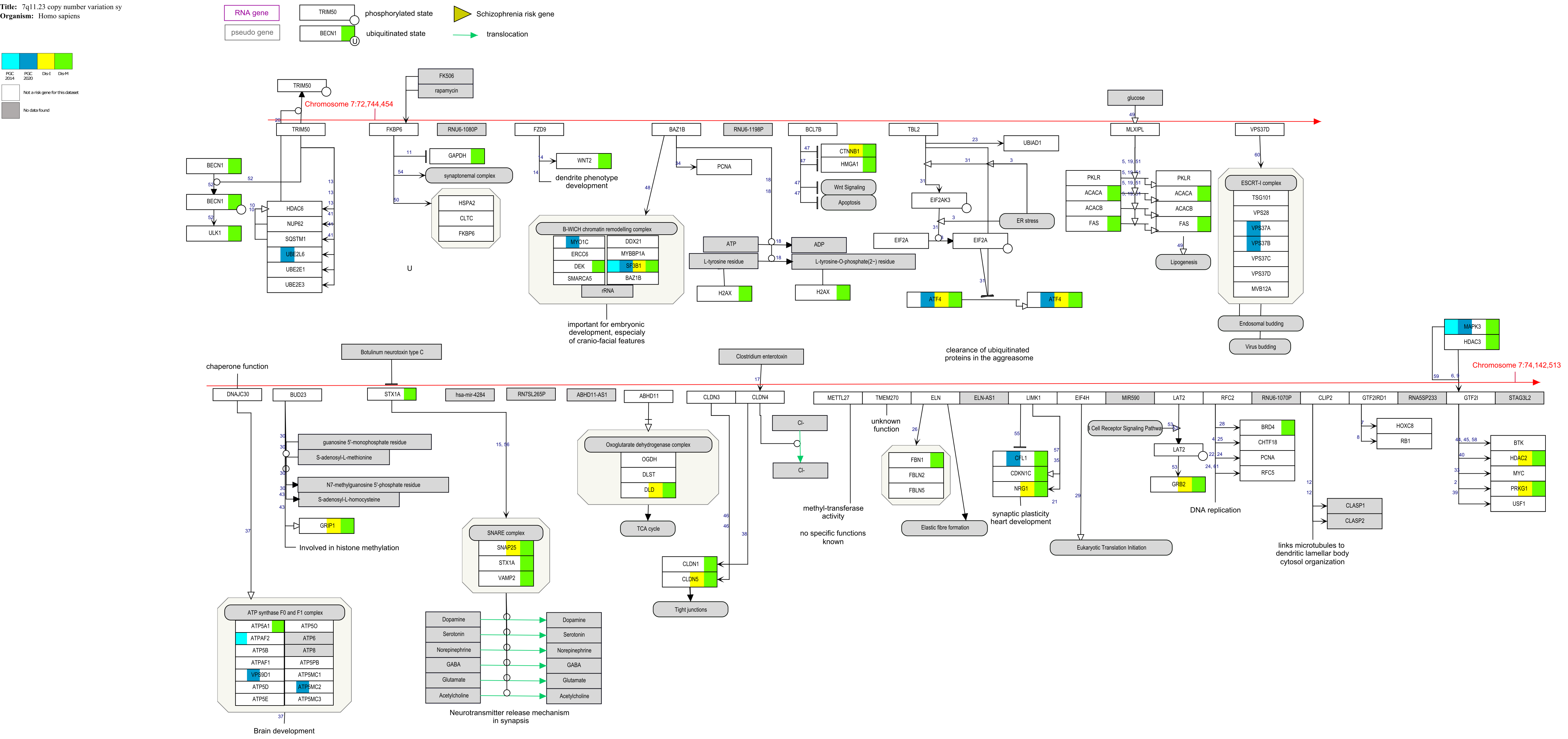

**Title:** 15q11.2 copy number variation sy
**Organism:** Homo sapiens

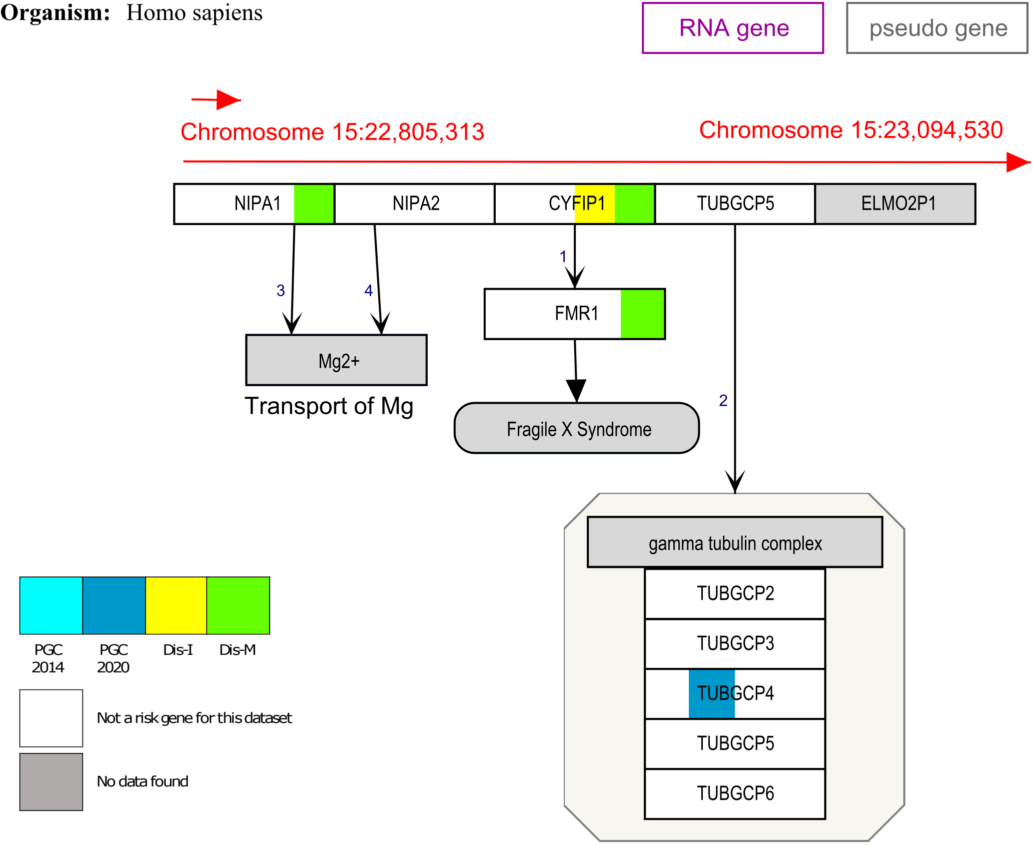

**Title:** 15q13.3 copy number variation sy
**Organism:** Homo sapiens

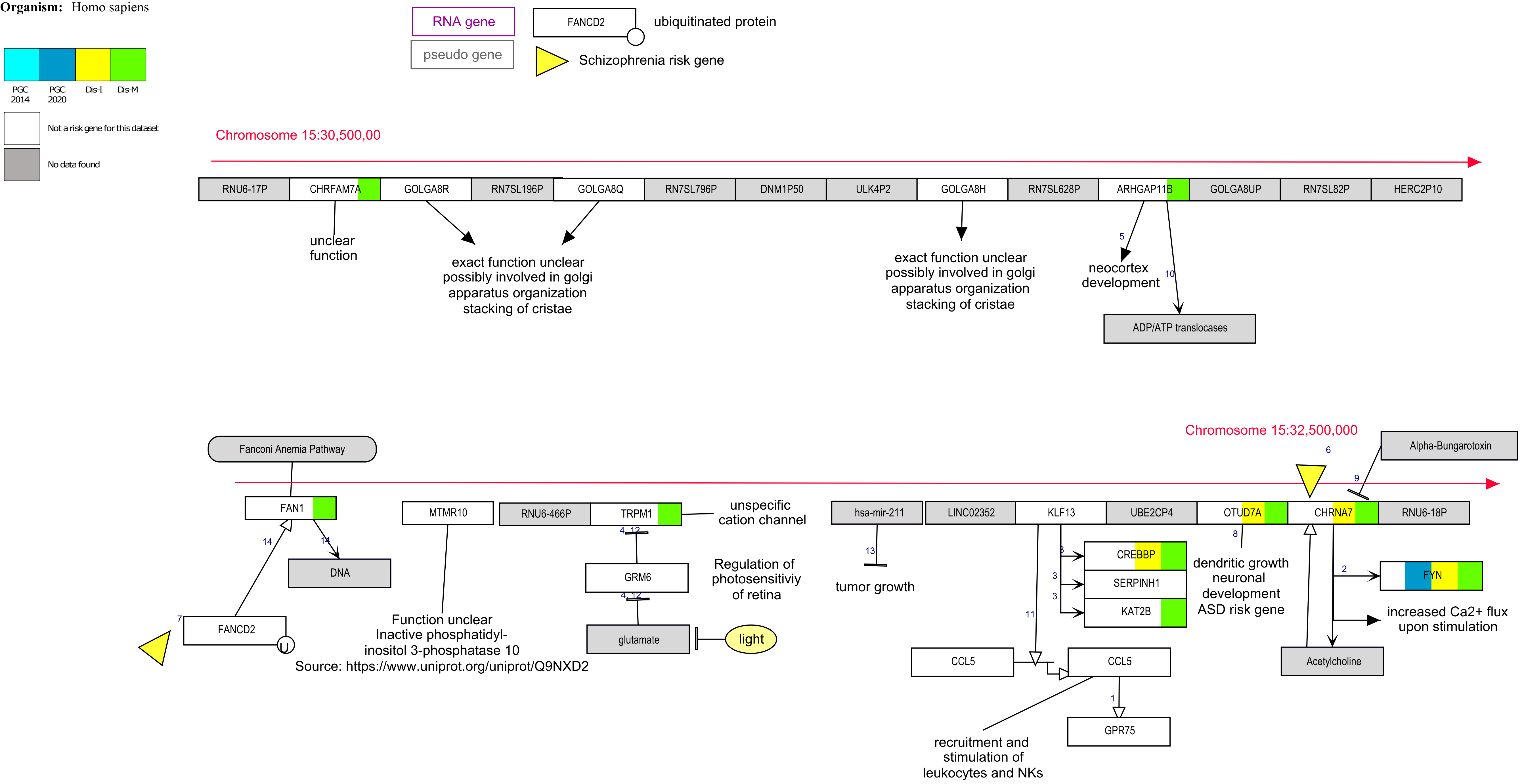

**Title:** 16p11.2 distal deletion syndrome
**Organism:** Homo sapiens

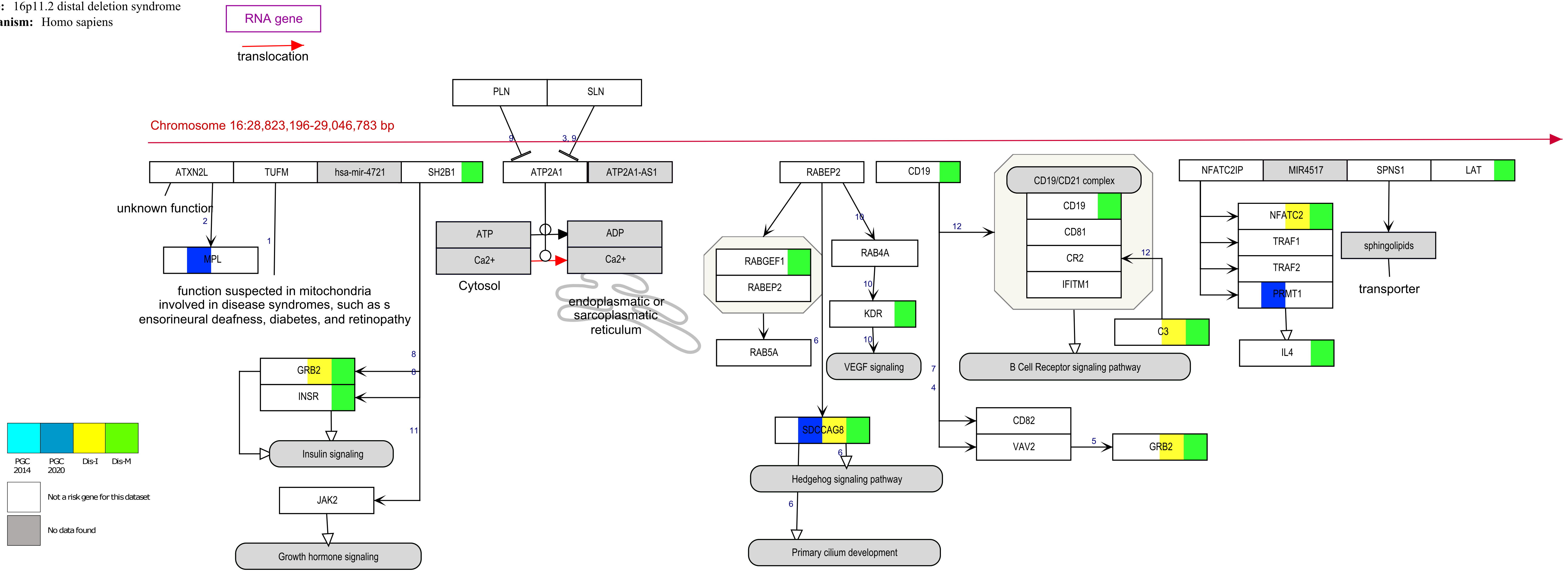

**Title:** 16p11.2 proximal deletion syndro[illegible]
**Organism:** Homo sapiens

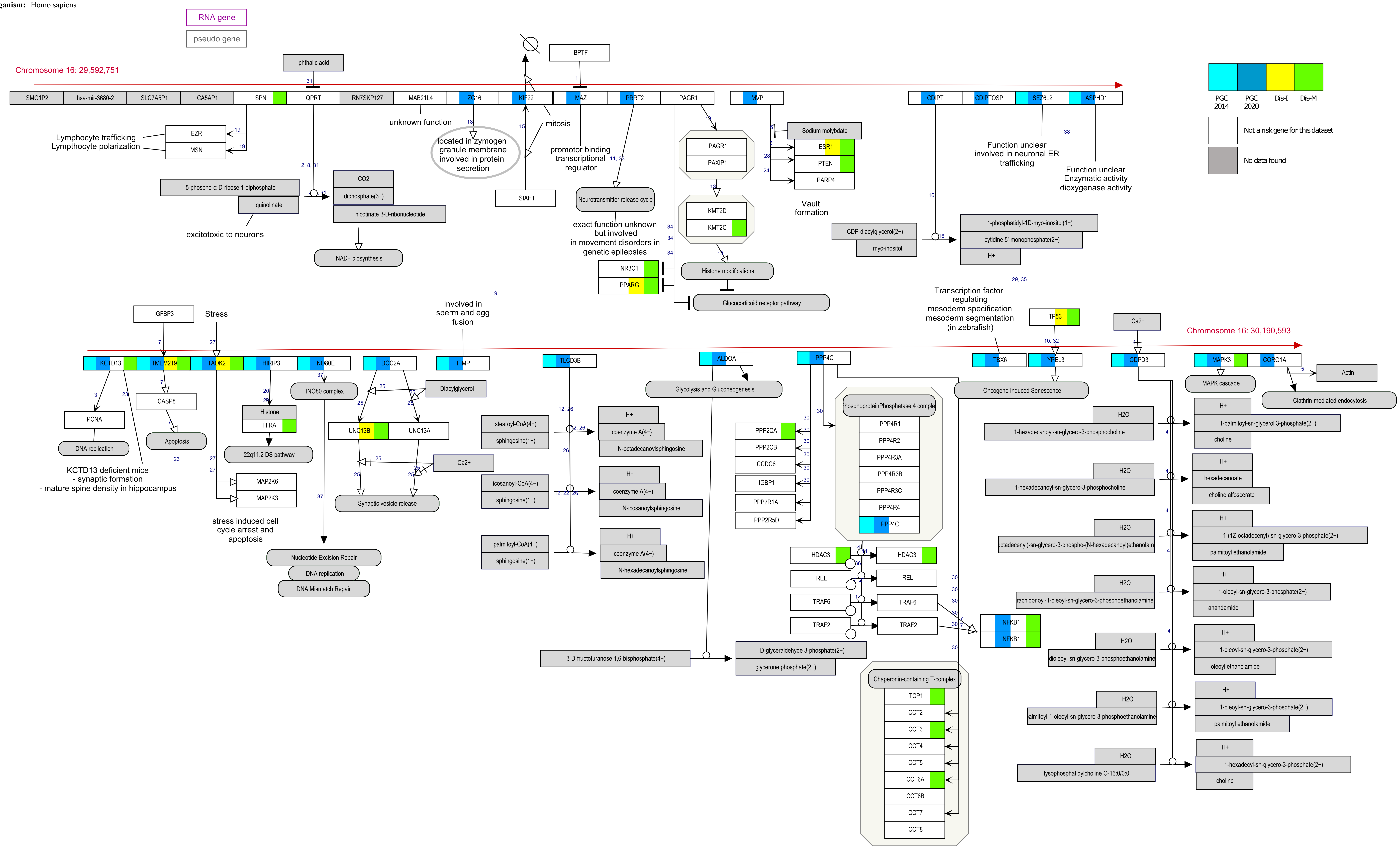

**Title:** 22q11.2 copy number variation sy
**Organism:** Homo sapiens

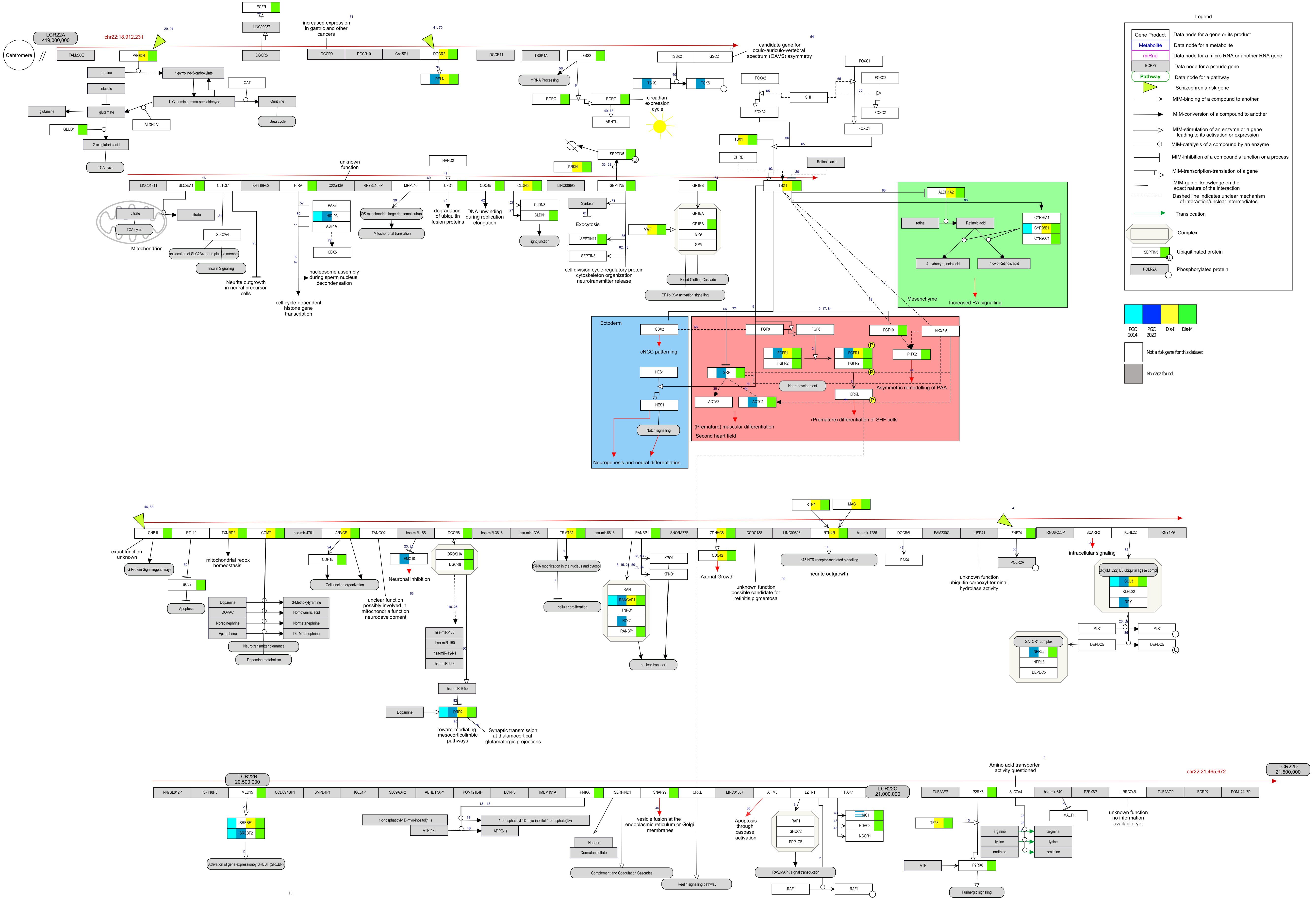
