## Supplementary information 1 for "Converging pathways found in copy number variation syndromes with high schizophrenia risk"

**Supplementary information 1 - Search strategy**

To identify the molecular interaction partners of a gene, gene product (RNA or protein) the following resources were used:

PubMed [https://pubmed.ncbi.nlm.nih.gov/]: The HGNC symbol, and/or the gene name were queried and literature was selected that focusses on the elucidation of the molecular function or the molecular interactions of this gene product. The PubMed identifier of the publication was added to the pathway as reference for this information.

UniProt [https://www.uniprot.org/]: After querying the HGNC symbol or the gene name the human protein with the curation level “reviewed” was selected. The Function section of the protein display page usually gives a short introduction to the most important protein functions and the Interaction section shows the known protein binding partners. STRING [https://string-db.org/] is one of the databases from which the protein binding information is derived from. For both information literature evidence is usually given in UniProt database. These references were added to the pathway. If the protein has a catalytic function, there is usually the known or the most important catalytic reactions described. These were annotated with a RHEA identifier and the chemical compounds were added using ChEBI identifiers.
