## Supplementary information 2 for "Converging pathways found in copy number variation syndromes with high schizophrenia risk"

**Supplementary material 2 – Permutation statistics**

The code for permutation statistics is available on <https://github.com/fehrhart/code/blob/master/R_permutationtest/R_CNVpathways>

1. **Eight CNV pathways vs. BDNF pathway (WP2380)**

- The genes in the eight CNV pathways were replaced by a random selection of 13562 genes (genes available in WikiPathways). The number of genes per pathway remained the same. Each gene can occur only once per CNV pathway.
- The genes in the BDNF pathway remained the same.
- The number of shared genes between the random selection and the BDNF pathway was counted.

| **Pathway**  *Number of genes* | **WP4940**  **15q11.2**  *9* | **WP4942**  **15q13.3**  *22* | **WP4905**  **1q21.1**  *28* | **WP4950**  **16p11.2 distal**  *33* |
| --- | --- | --- | --- | --- |
| Mean and sd | 0.09 ± 0.31 | 0.24 ± 0.48 | 0.28 ± 0.52 | 0.35 ± 0.59 |
| Plot | 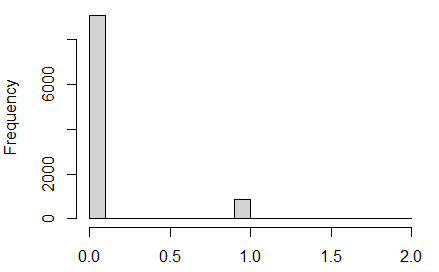 | 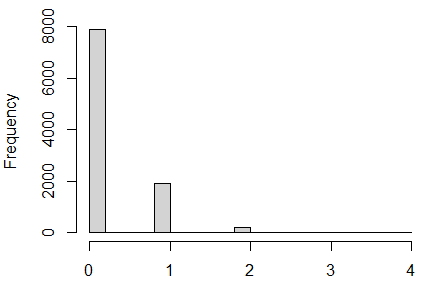 | 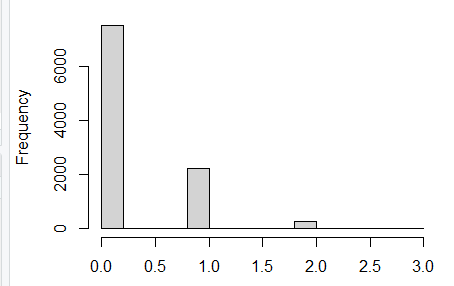 | 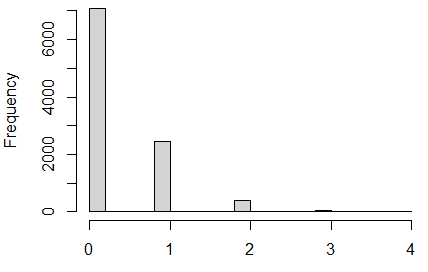 |
| **Pathway**  *Number of genes* | **WP4906**  **3q29**  *73* | **WP4949**  **16p11.2 proximal**  *79* | **WP4932**  **7q11.23**  *107* | **WP4657**  **22q11.2**  *140* |
| Mean and sd | 0.78 ± 0.89 | 0.82 ± 0.90 | 1.13 ± 1.06 | 1.47 ± 1.21 |
| Plot | 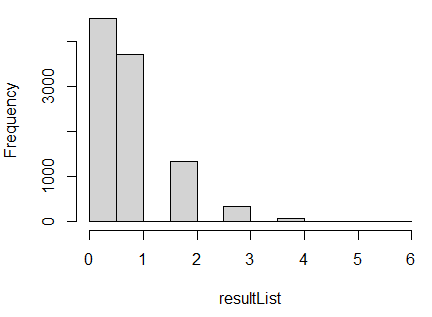 | 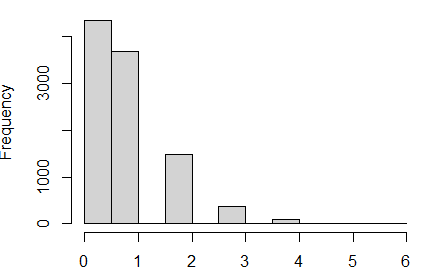 | 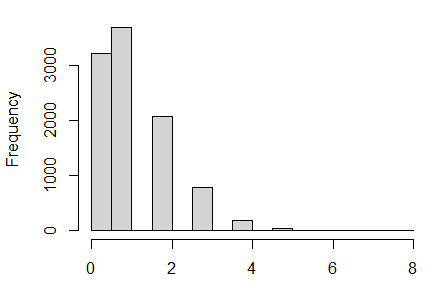 | 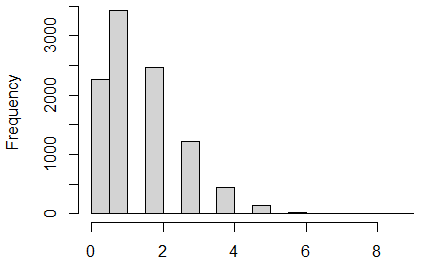 |

**Supplementary material 2 - Table 1:** The table shows the average number (and standard deviation) of genes from a randomly selected gene set of the same size as the respective CNV pathway overlapping with the BDNF pathway vs. the frequency of the event. The plots are shown for **each individual** of the eight CNV pathways (number of permutations =10 000).

Average number of overlapping gens with BDNF pathway

| number of genes in the pathway | mean |
| --- | --- |
| 9 | 0.09 |
| 22 | 0.24 |
| 28 | 0.28 |
| 33 | 0.35 |
| 73 | 0.78 |
| 79 | 0.82 |
| 107 | 1.13 |
| 140 | 1.47 |

**Supplementary material 2 - Figure 1:** Plot of mean and standard deviation from the table above of shared genes from random samples with the BDNF pathway vs. pathway size.

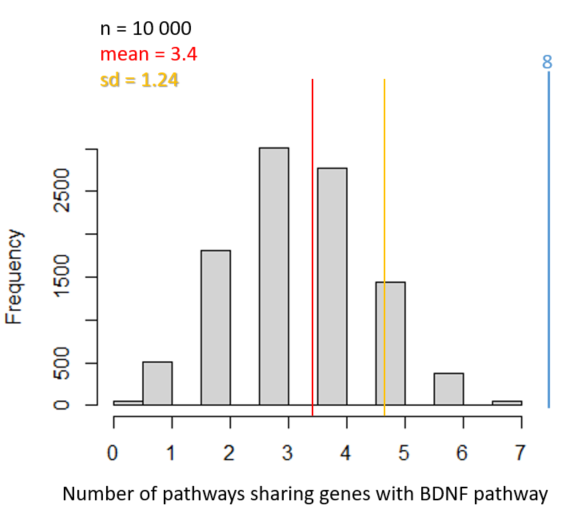

**Supplementary material 2 - Figure 2:** Number of pathways sharing genes with the BDNF pathway per permutation round. The red line indicates the mean number, the yellow line the standard deviation. In this simulation, in none of the 10 000 runs all eight CNV pathways shared genes with the BDNF pathway (blue line).

**Conclusion:** The probability of a randomly selected gene set to share genes with the BDNF pathway is depending on the size of the randomly selected gene set (the pathway) The larger the pathway, the higher the probability of overlap. Pathways with more than 100 genes are likely to share a (one) random gene with the BDNF pathway. However, the probability that all eight CNV pathways share genes with the BDNF pathway by chance is extremely low (Figure 2).

2. **Seven CNV pathways vs. the other seven overlapping pathways**

**Supplementary material 2 - Table 2:** In contrast to Table 1, here the number of CNV pathways (7 – without the smallest CNV pathway WP4940) that share genes with the respective pathway in the table is shown (similar to Figure 2). Mean and sd indicate the average number of overlapping CNV pathways (= random gene sets of the size of the pathway) with the respective pathway in the table.

| **Pathway**  *Number of genes* | **WP4352**  **Ciliary landscape**  *213* | **WP2034**  **Leptin signaling**  *75* | **WP3888**  **VEGFA-VEGFR2 signaling**  *431* | **WP2036**  **TWEAK signaling**  *42* |
| --- | --- | --- | --- | --- |
| Mean and sd | 4.2 ± 1.2 | 2.1 ± 1.1 | 5.5 ± 1.0 | 1.3 ± 1.0 |
| Plot | 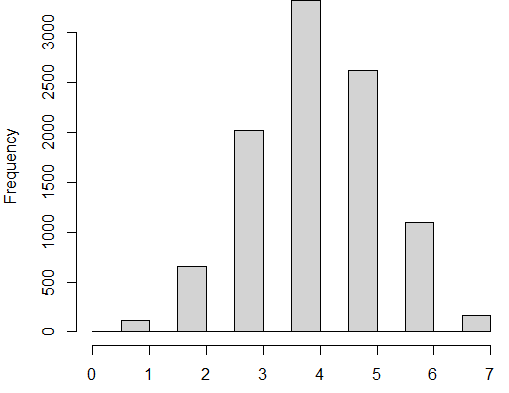 | 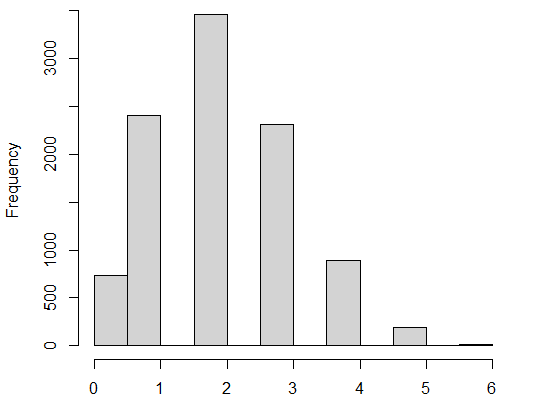 | 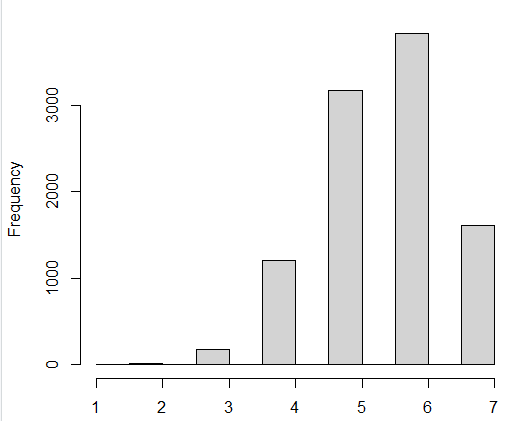 | 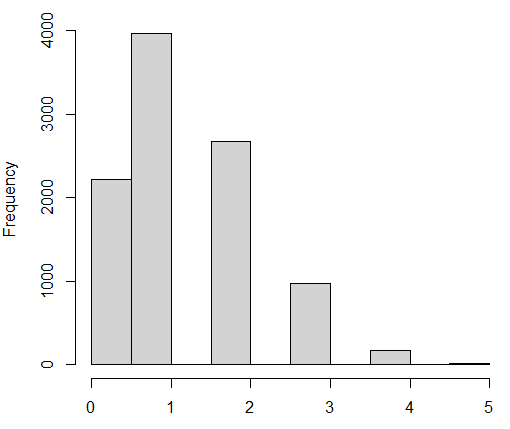 |
| **Pathway**  *Number of genes* | **WP4754**  **IL-18 signaling**  *273* | **WP306**  **Focal adhesion**  *198* | **WP4659**  **Gastrin signaling**  *114* |  |
| Mean and sd | 4.7 ± 1.1 | 4.0 ± 1.2 | 2.9 ± 1.2 |  |
| Plot | 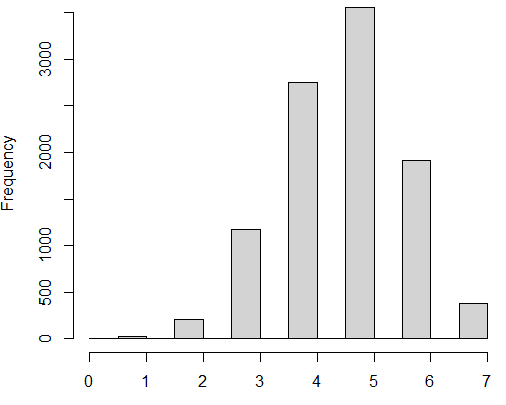 | 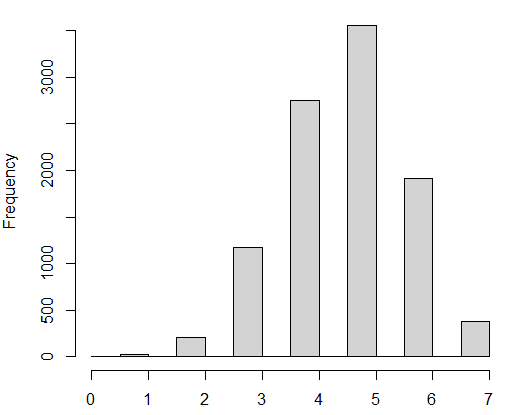 | 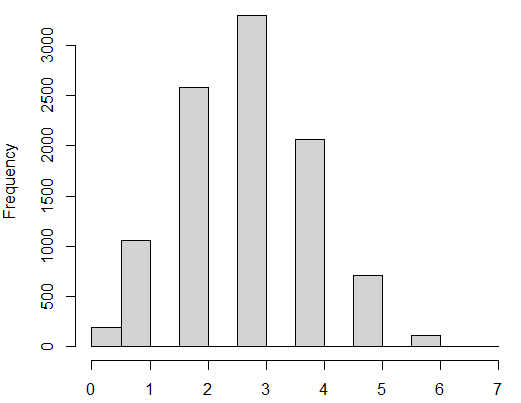 |  |
