## Supplementary overlapping pathways figures for "Converging pathways found in copy number variation syndromes with high schizophrenia risk"

**Title:** Ciliary landscape
**Organism:** Homo sapiens

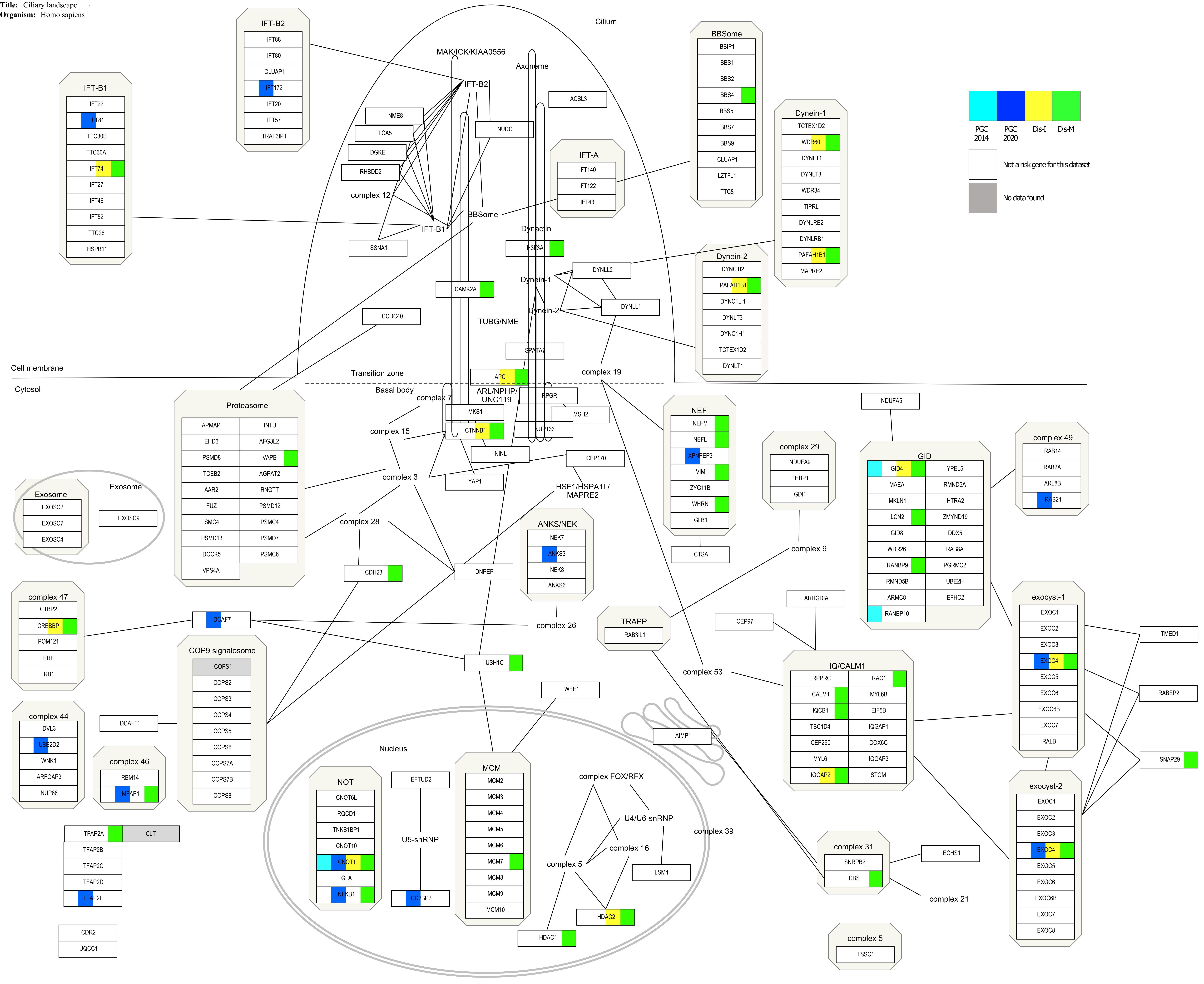

**Title:** Focal adhesion 1-17
**Organism:** Homo sapiens

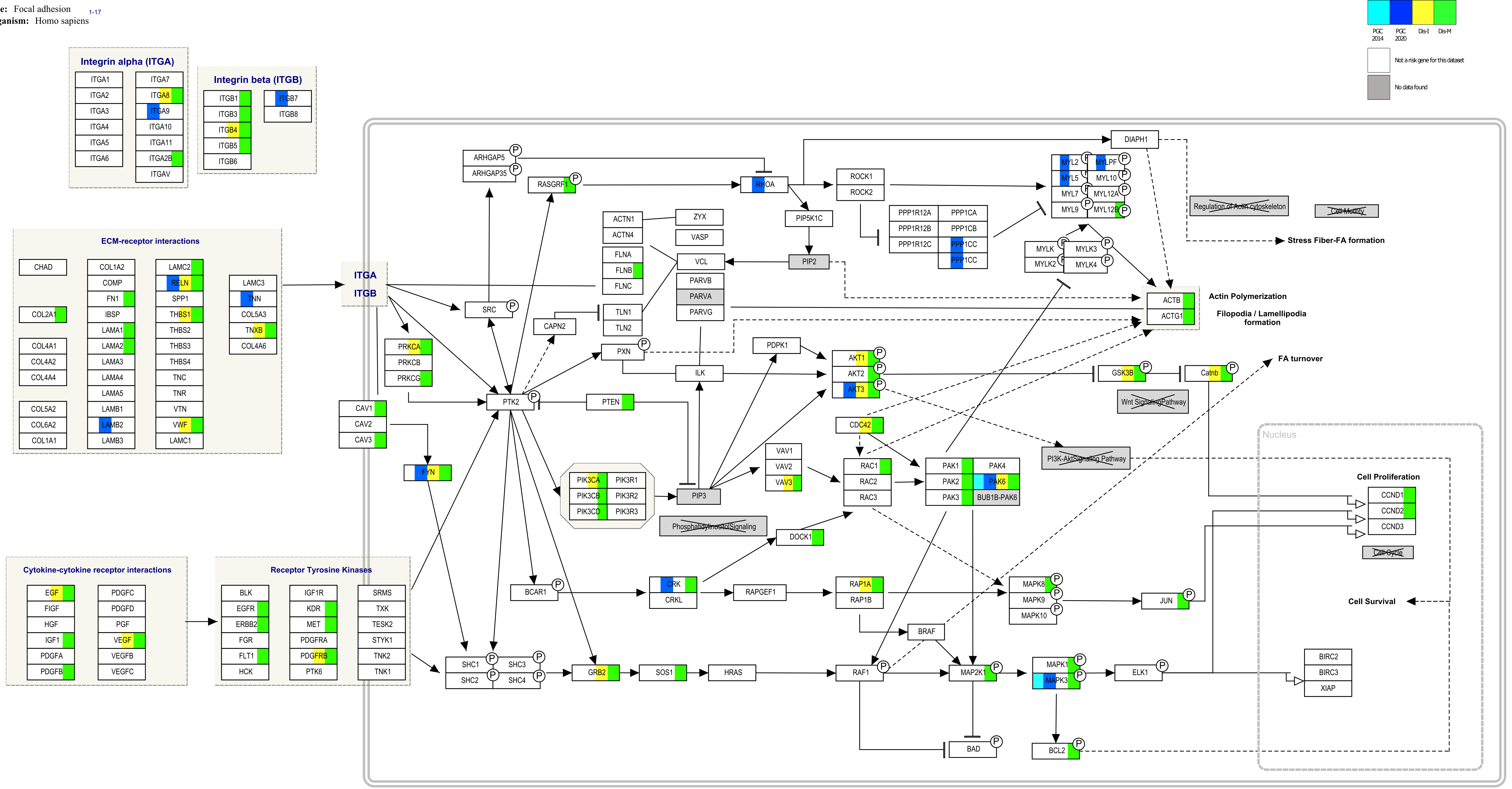

**Title:** Gastrin signaling pathway
**Maintained by:** Institute of Bioinformat
**Availability:** Creative Commons Licens
**Last modified:** 06-SEP-2013
**Organism:** Homo sapiens
**Data Source:** NetPath

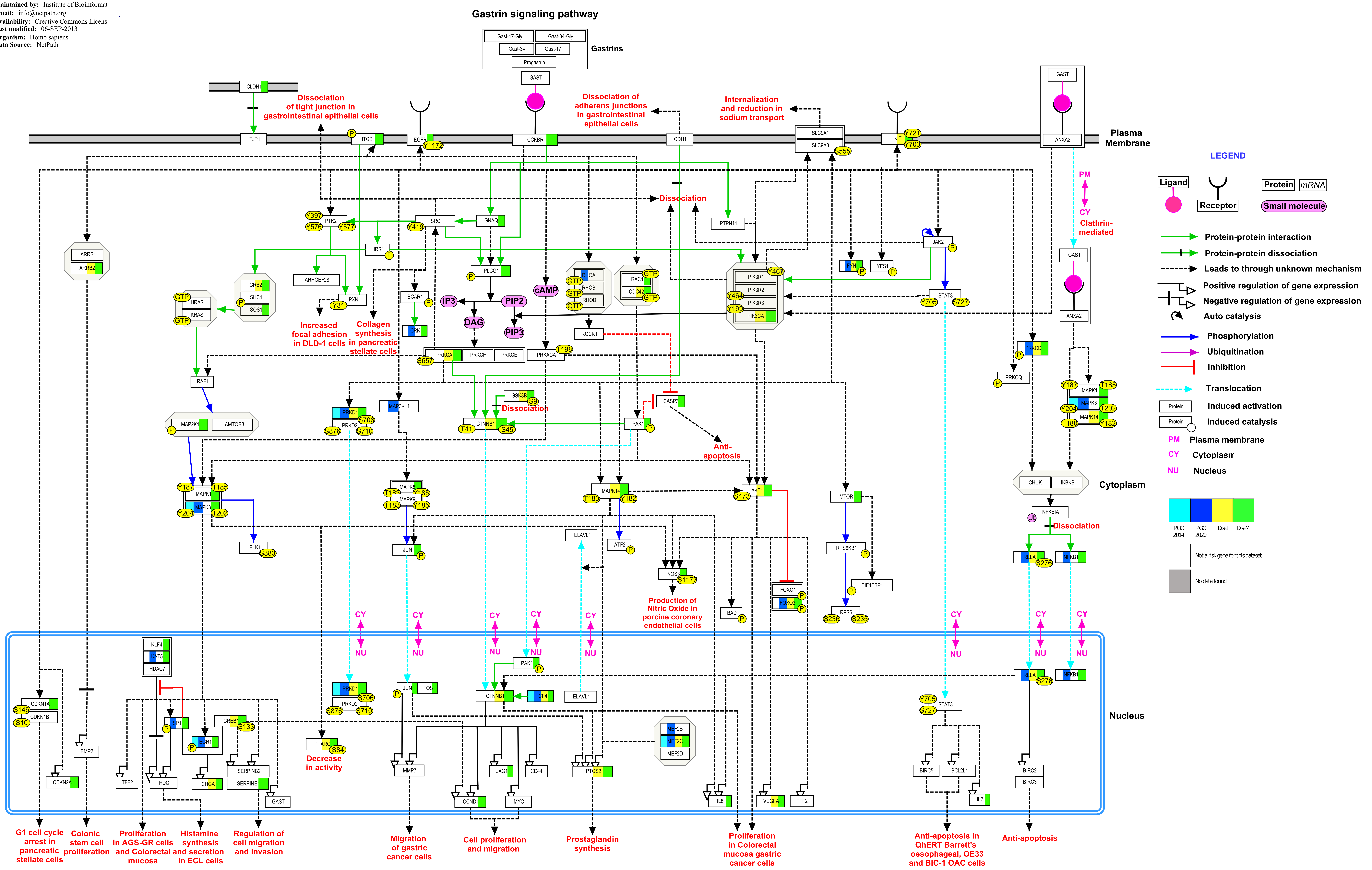

**Title:** IL-18 signaling pathway
**Availability:** Freely available under crea
**Last modified:** November 11, 2019
**Organism:** Homo sapiens

1

### IL-18 Signaling Pathway

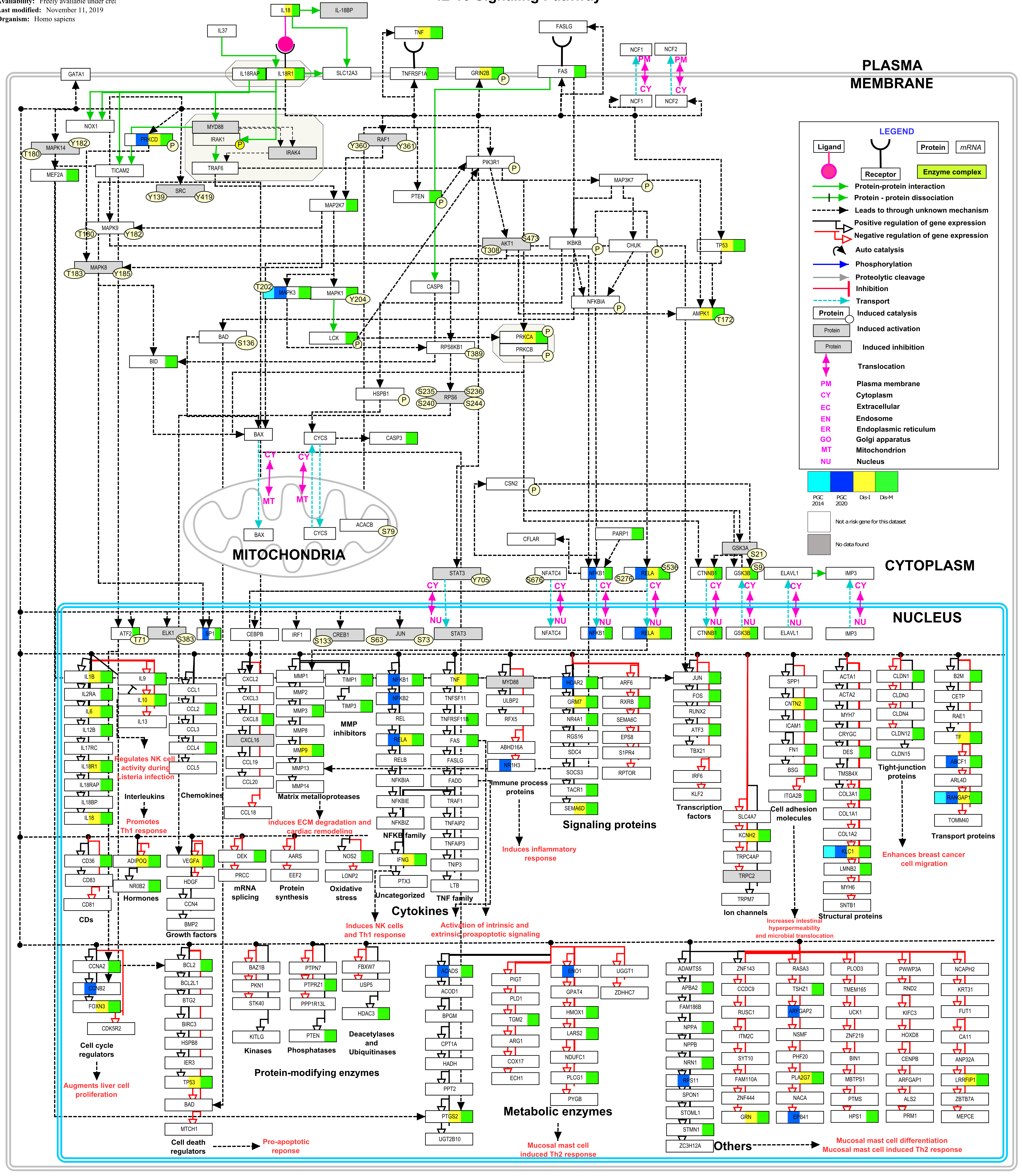

**Title:** Leptin signaling pathway
**Availability:** Freely available under con
**Organism:** Homo sapiens

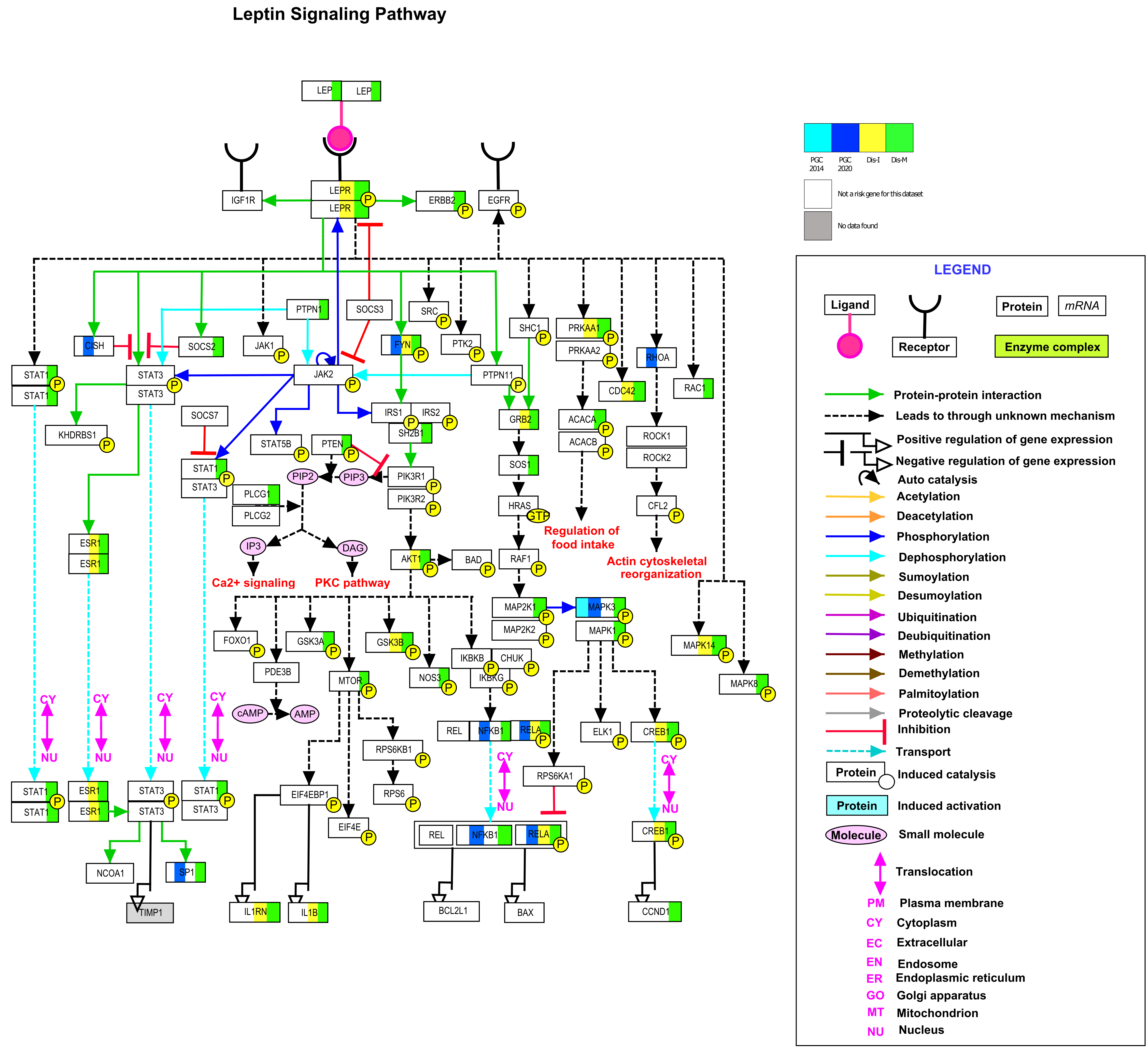

**Title:** TNF-related weak inducer of apop
**Availability:** Freely available under Cre
**Organism:** Homo sapiens

1

#### TWEAK Signaling Pathway

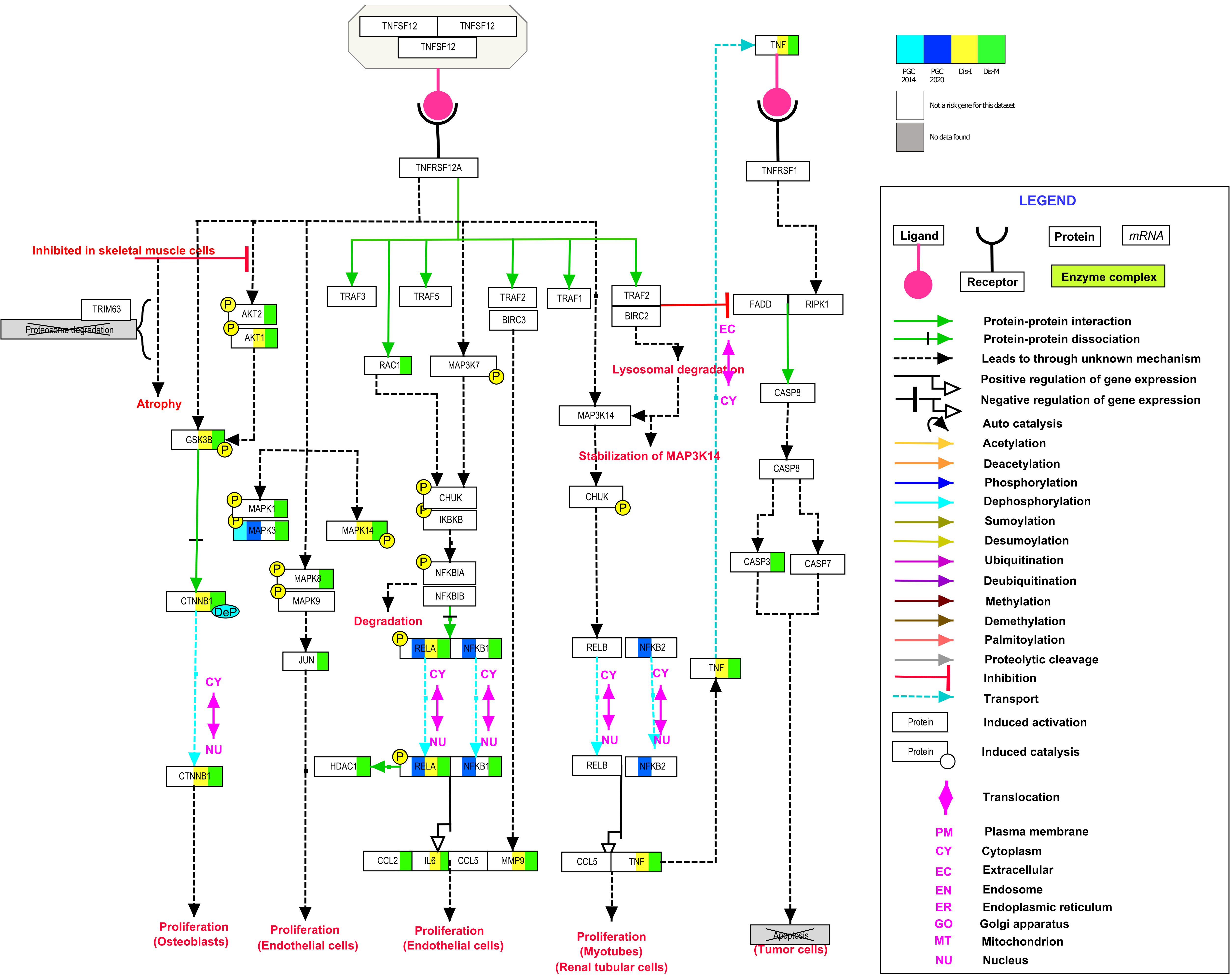

**Title:** VEGFA-VEGFR2 signaling pathw[illegible]
**Maintained by:** Dr. Rajesh Raju, Prof. [illegible]
**Availability:** Creative Commons Licens[illegible]
**Organism:** Homo sapiens
**Data Source:** PRS Lab

### VEGFA SIGNALING PATHWAY IN ENDOTHELIAL CELLS

| | | | | |
|---|---|---|---|---|
| ABCF2 | CNP | USP10 | MYH11 | SSR3 |
| ACOT9 | CRIP2 | FAM120A | MYO1C | SET |
| ACTG1 | CSRP1 | YWHAE | MYO6 | SPIRE1 |
| AKAP2 | CSRP2 | FGD5 | NAP1L1 | SRP54 |
| ALDOA | CYR61 | ZC3H15 | NUMB | SRPK1 |
| AP2A1 | DECR1 | FHOD1 | MOV10 | SSR4 |
| AP2S1 | DHX36 | VPS39 | NAPA | STIP1 |
| ASCC3 | DNAJA1 | FLII | NCL | SYNJ1 |
| ALB | DNAJB4 | FMNL3 | NEXN | TAOK2 |
| ARF4 | DPM1 | GAPDH | OCRL | TBCA |
| ARMCX1 | EIF3H | FARSB | P4HA2 | TCEB1 |
| ARPC5L | DHX29 | FN1 | P4HB | TFCP2 |
| ANXA1 | DSC1 | FSCN1 | PABPC1 | TKT |
| TUBB8 | EIF3D | FXR2 | PBXIP1 | TMOD1 |
| BSG | EIF3F | GIGYF2 | PDE4DIP | TMOD3 |
| CALR | EIF4G1 | GLUD1 | PFN1 | TPP1 |
| CAPZB | EIF4G2 | GRSF1 | PGD | TFAM |
| CCT7 | EPB41 | HBD | PGK1 | TMSB10 |
| SHROOM2 | TXNDC5 | COPG1 | PLOD3 | TMSB4X |
| CALU | EPRS | HMGB1 | PRDX2 | TPM3 |
| CGNL1 | UBAP2L | HSP90AB2P | PDIA6 | TRIP4 |
| CLIC1 | EWSR1 | HTRA1 | PPM1G | TUBA1C |
| PPP2CA | PRDX6 | IQGAP1 | KANK1 | LDHA |
| QKI | RBM39 | RCN1 | RPL10A | RPL27 |
| PRKRA | PRRC2C | PSMD11 | PSMD4 | PTMA |
| INPP5K | LMAN1 | LRRC59 | LRRFIP2 | LUC7L |
| HYOU1 | IDH2 | PPP1CA | SND1 | MICAL2 |
| RPL7 | RPLP2 | RPS11 | RPL39P5 | RPL5 |
| PTPN14 | RPL13A | RPL18A | RPL26 | SARS |
| ATP6V0D1 | HSP90AA1 | KATNAL2 | SH3BGRL3 | |
| SLC25A11 | ATP6V1E1 | CACNA2D1 | CCDC124 | |

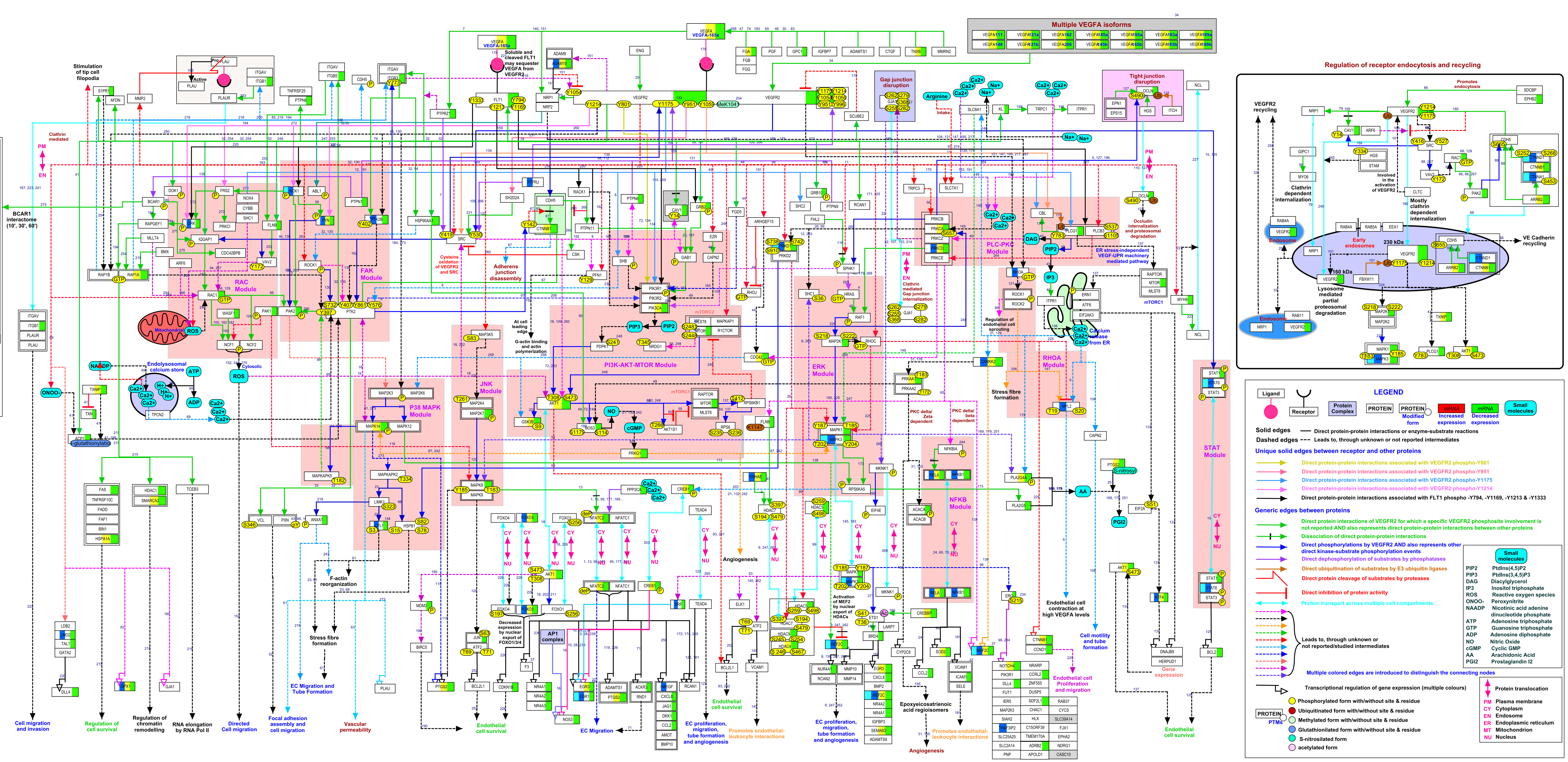

**Title:** Brain-derived neurotrophic factor
**Maintained by:** IOB, Pandey Lab
**Availability:** Adaptive commons creativ
**Last modified:** 25-NOV-2012
**Organism:** Homo sapiens

PGC 2014 | PGC 2020 | Dis-I | Dis-M

Not a risk gene for this dataset

No data found

### Brain-derived neurotrophic factor signaling pathway

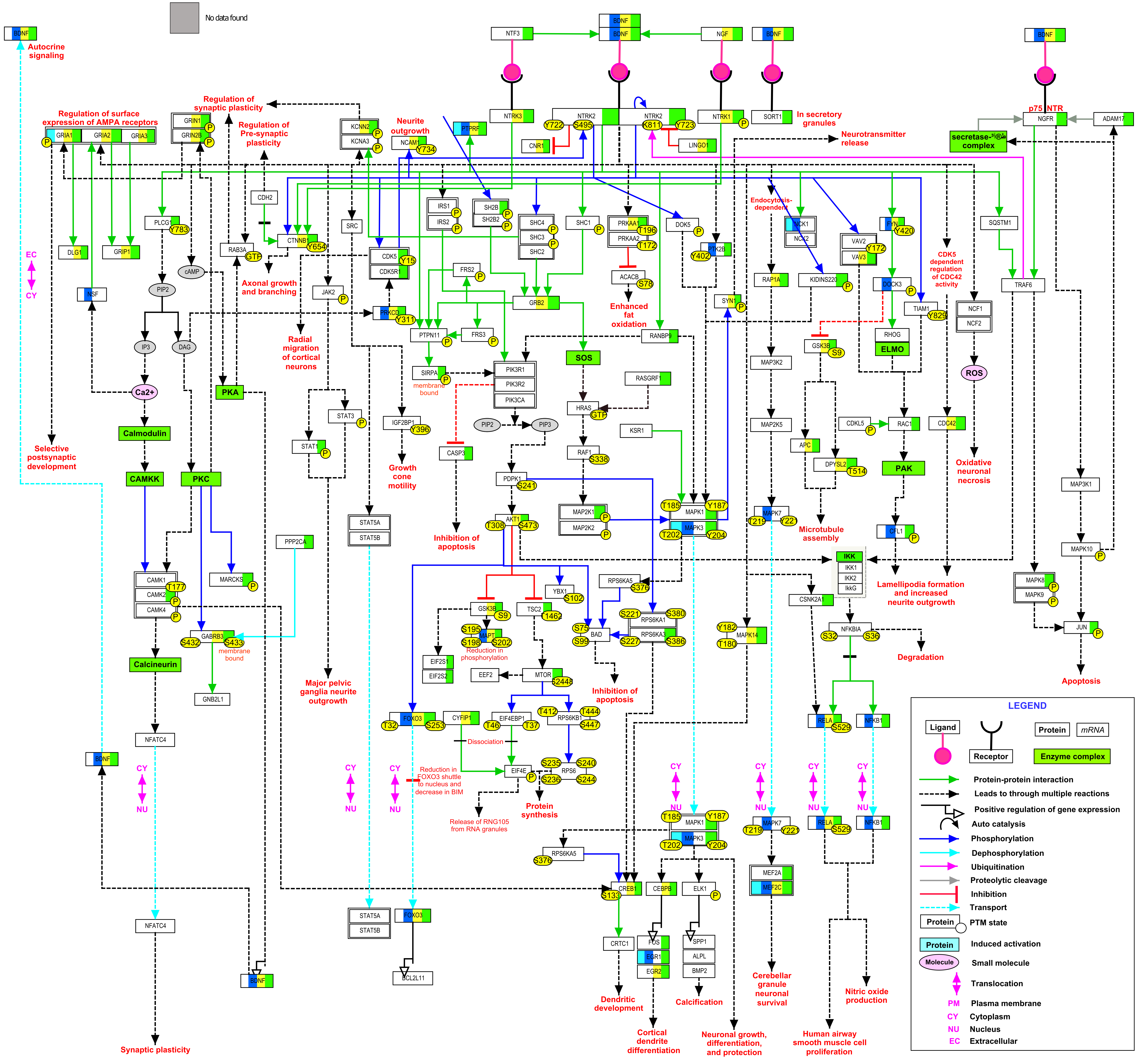
